# Development of CRISPR as a prophylactic strategy to combat novel coronavirus and influenza

**DOI:** 10.1101/2020.03.13.991307

**Authors:** Timothy R. Abbott, Girija Dhamdhere, Yanxia Liu, Xueqiu Lin, Laine Goudy, Leiping Zeng, Augustine Chemparathy, Stephen Chmura, Nicholas S. Heaton, Robert Debs, Tara Pande, Drew Endy, Marie La Russa, David B. Lewis, Lei S. Qi

## Abstract

The outbreak of the coronavirus disease 2019 (COVID-19), caused by the Severe Acute Respiratory Syndrome coronavirus 2 (SARS-CoV-2), has infected more than 100,000 people worldwide with over 3,000 deaths since December 2019. There is no cure for COVID-19 and the vaccine development is estimated to require 12-18 months. Here we demonstrate a CRISPR-Cas13-based strategy, PAC-MAN (<u>P</u>rophylactic <u>A</u>ntiviral <u>C</u>RISPR in huMAN cells), for viral inhibition that can effectively degrade SARS-CoV-2 sequences and live influenza A virus (IAV) genome in human lung epithelial cells. We designed and screened a group of CRISPR RNAs (crRNAs) targeting conserved viral regions and identified functional crRNAs for cleaving SARS-CoV-2. The approach is effective in reducing respiratory cell viral replication for H1N1 IAV. Our bioinformatic analysis showed a group of only six crRNAs can target more than 90% of all coronaviruses. The PAC-MAN approach is potentially a rapidly implementable pan-coronavirus strategy to deal with emerging pandemic strains.

## INTRODUCTION

The world is currently faced with a pandemic of novel coronavirus disease 2019 (COVID-19), which is caused by Severe Acute Respiratory Syndrome coronavirus 2 (SARS-CoV-2) and has no vaccine or cure. It is predicted the development of a safe and effective vaccine to prevent COVID-19 will take one year to 18 months, by which time it is likely that several hundreds of thousands to millions of people may have been infected. With a rapidly growing number of cases and deaths around the world, this emerging threat requires a nimble and targeted means of protection. Since coronaviruses causing COVID-19, Severe Acute Respiratory Syndrome (SARS), and Middle East Respiratory Syndrome (MERS) are able to suddenly transfer to humans from diverse animal hosts that act as viral reservoirs, there is a pressing need to develop methods to combat other potential coronaviruses that may emerge in the future^1-3^. A recent report further showed two strains (L and S) of SARS-CoV-2 with different genome sequences are circulating and likely evolving, further highlighting the need for a pan-coronavirus vaccination strategy^4^.

The novel coronavirus causing COVID-19 belongs to a family of positive-sense RNA viruses, which typically infect the upper and lower respiratory tracks and cause disease by direct cytotoxic effects and the induction of host cytokines disease ^5^. The SARS-CoV-2 life cycle is likely similar to closely related coronaviruses such as the virus that causes SARS, in which the virus enters the cell, releases its RNA genome into the cytoplasm, and synthesizes negative-sense genomic and subgenomic RNAs from which viral mRNAs and a new copy of the positive sense viral genome are synthesized (**Fig. 1A**)^6,7^. Whereas traditional vaccines work by priming the human immune system to recognize viral proteins or weakened viruses and decrease viral entry into cells^8^, the alternative antiviral approach we propose here relies on a CRISPR-based system for recognizing and degrading the intracellular viral genome and its resulting viral mRNAs (**Fig. 1B**). Targeting the positive-sense genome and viral mRNAs to simultaneously degrade viral genome templates for replication and viral gene expression would be expected to robustly limit viral replication.

To inhibit RNA viruses in human cells, we used the CRISPR-Cas13d system derived from *Ruminococcus flavefaciens* XPD3002, a recently discovered RNA-guided RNA endonuclease^9,10^. Cas13d employs CRISPR-associated RNAs (crRNAs) that contain a customizable 22 nucleotide (nt) spacer sequence that can direct the Cas13d protein to specific RNA molecules for targeted RNA degradation. The high catalytic activity of Cas13d in mammalian cells provides a potential mechanism for targeting SARS-CoV-2 for specific viral RNA genome degradation and viral gene expression inhibition. Because of its small size (967 amino acids), high specificity, and strong catalytic activity, we chose Cas13d rather than other Cas13 proteins to target and destroy RNA viruses including SARS-CoV-2 and Influenza A virus (IAV)^10-12^.

In this work, we developed a <u>P</u>rophylactic <u>A</u>ntiviral <u>C</u>RISPR in hu<u>MAN</u> cells (**PAC-MAN**) strategy as a form of genetic intervention to target SARS-CoV-2, IAV, and potentially all sequenced coronaviruses. We created a bioinformatic pipeline to define conserved regions across the SARS-CoV-2 genomes and target these regions using CRISPR-Cas13d for viral genome degradation and viral gene inhibition. At the time of paper submission, there is no widely available laboratory strains of SARS-CoV-2. Therefore, we tested our approach using synthesized fragments of SARS-CoV-2, as well as with live H1N1 IAV. We designed and screened a panel of crRNA pools targeting conserved viral regions and defined the most effective crRNAs. We demonstrated the ability of our approach to cleave SARS-CoV-2 fragments and to reduce the replication of IAV in human lung epithelial cells. Our bioinformatics analysis revealed a group of six crRNAs that can target 91% of sequenced coronaviruses, as well as a group of 22 crRNAs able to target all sequenced coronaviruses. The potential pan-coronavirus protection using CRISPR-Cas13d offers an alternative and complementary approach over traditional pharmaceuticals or vaccines. Through the use of crRNA pools targeting different regions of the same virus or different strains of coronavirus, this system could possibly buffer against viral evolution and escape, as well as enable rapid development and deployment against emerging viruses. When combined with an effective delivery platform, PAC-MAN is a promising strategy to combat not only coronaviruses including that causing COVID-19, but also a broad range of other genera of viruses.

**Figure 1.**
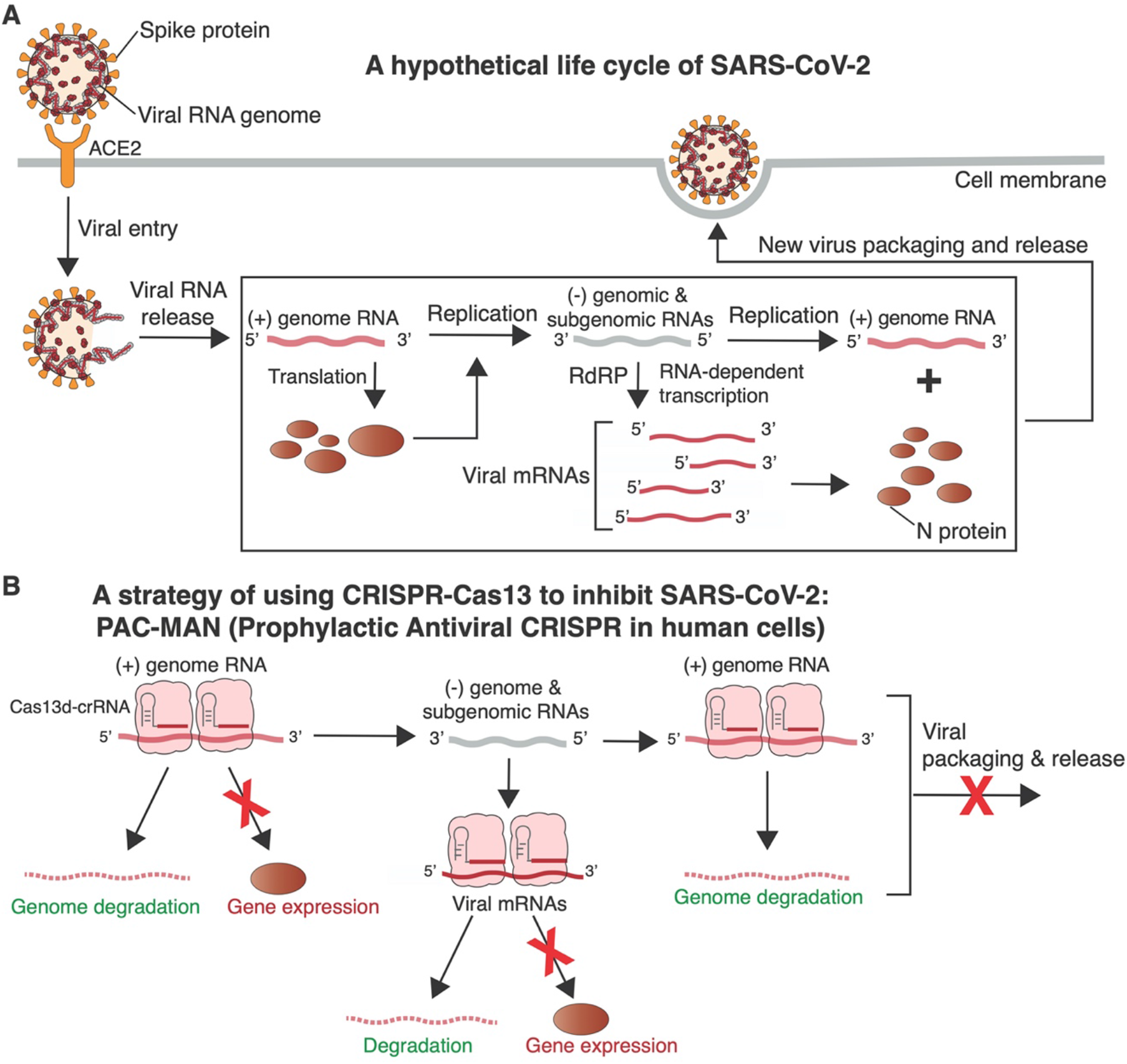
The hypothetical life cycle of SARS-CoV-2 and the PAC-MAC approach for anti-COVID-19. **(A)** A hypothetical life cycle of SARS-CoV-2 based on what is known about other coronavirus life cycles. SARS-CoV-2 virions bind to the ACE2 receptor on the surface of cells via interactions with the Spike protein. Upon viral release, the positive strand RNA genome serves as a template to make negative strand genomic and subgenomic templates, which are used to produce more copies of the positive strand viral genome and viral mRNAs. **(B)** Cas13d can inhibit viral function and replication by directly targeting and cleaving all viral positive-sense RNA.

## RESULTS

### Computational design and analysis of targetable regions on SARS-CoV-2

To create effective and specific crRNA sequences to target and cleave SARS-CoV-2, we first performed a bioinformatic analysis by aligning published SARS-CoV-2 genomes from 47 patients with SARS-CoV and MERS-CoV genomes at the single-nucleotide resolution level. SARS-CoV-2 has a single-stranded RNA genome with ~30,000 nucleotides that encodes 12 putative, functional open reading frames^13-15^. Our analysis found regions with a high conservation between 47 SARS-CoV-2, SARS-CoV and MERS-CoV genomes (**Fig. 2A**). The two highly-conserved regions respectively contain the RNA-dependent RNA polymerase (RdRP) gene in the polypeptide ORF1ab region, which maintains the proliferation of all coronaviruses, and the Nucleocapsid (N) gene at the 3’ end of the genome, which encodes the capsid protein for viral packaging.

**Figure 2.**
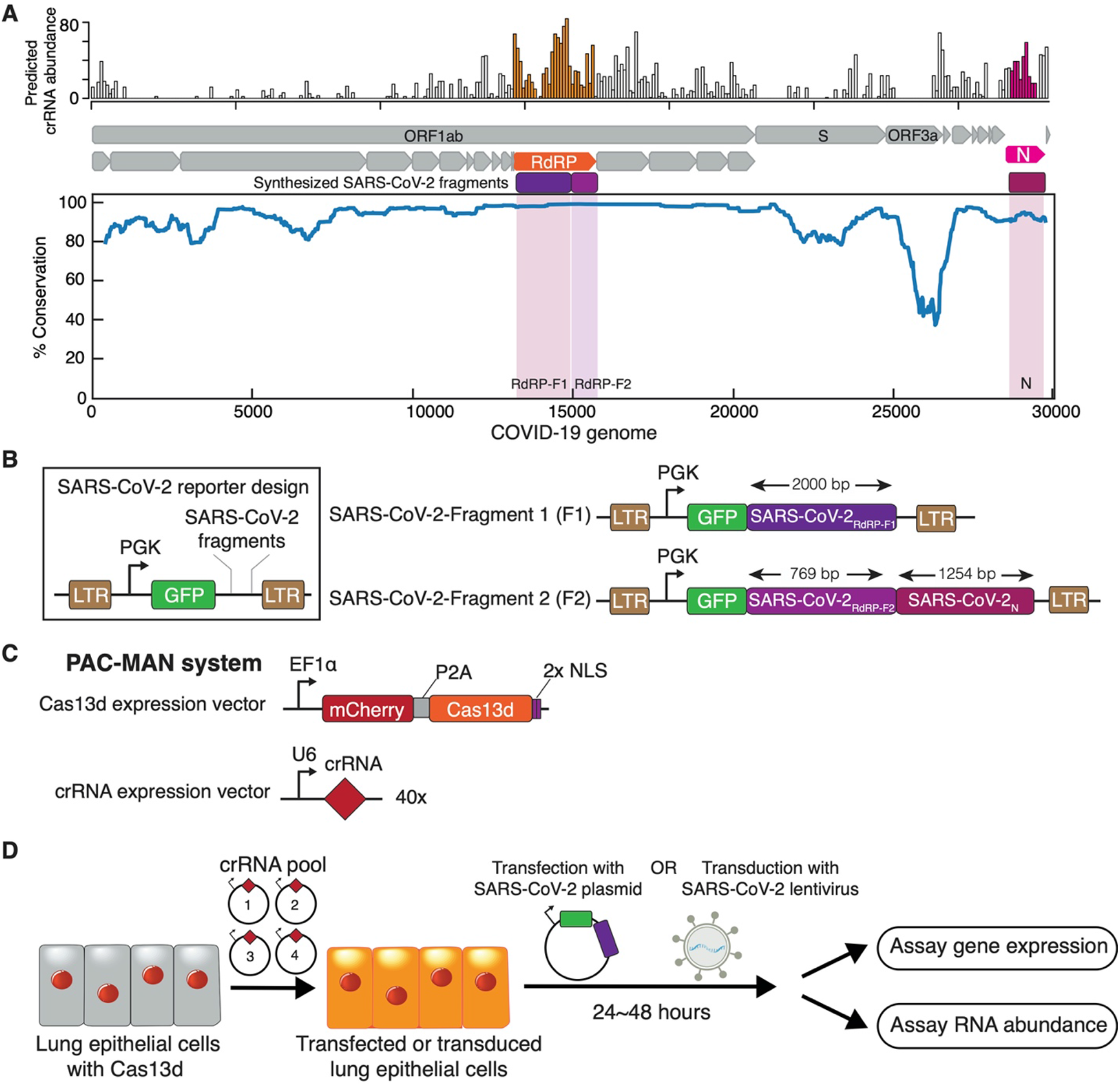
Bioinformatic analysis of CRISPR-Cas13d target sites for SARS-CoV-2 and the PAC-MAN system design. **(A)** Alignment of sequences from SARS-CoV-2 genomes from 47 patients along with SARS- and MERS-causing coronaviruses. Top: predicted abundance of crRNAs that are able to target SARS-CoV-2 genomes and SARS or MERS; Middle: annotation of genes in the SARS-CoV-2 genomes, along with conserved regions chosen to be synthesized into the SARS-CoV-2 reporters (in magenta and purple). Color regions indicate the synthesized SARS-CoV-2 fragments; Bottom: the percent of conservation between aligned viral genomes. See **Supplemental Table 2** for the designed crRNA sequences and **Supplemental Table 3** for synthesized SARS-CoV-2 fragments. **(B)** Schematic of the two reporters (SARS-CoV-2-F1/F2) created with synthesized viral sequences. SARS-CoV-2-F1 contains GFP fused to a portion of RdRP (RdRP-F1) and SARS-CoV-2-F2 contains GFP fused to portions of both RdRP (RDRP-F2) and N. **(C)** Schematics for the constructs used to express Cas13d or its crRNAs. **(D)** Workflow used to challenge Cas13d A549 lung epithelial cells with SARS-CoV-2 reporters.

To target highly conserved SARS-CoV-2 regions, we generated an *in silico* collection of all 3,802 possible crRNAs **(Supplemental Fig. 1).** After excluding crRNAs either predicted to have potential off-target binding (≤1 mismatch) in the human transcriptome or having poly-T (≥4 Ts) sequences that may prevent crRNA expression, we obtained a collection of 3,203 crRNAs (**Supplemental Table 1**). We designed and synthesized 40 crRNAs, with 20 crRNAs each targeting the conserved sequences of the RdRP and N genes (**Supplemental Table 2**). Our rationale for densely targeting these two regions was to significantly reduce both viral genomic RNA and mRNA templates for expressing essential viral proteins and to minimize the risk of loss of targeting activity by mutational escape.

### CRISPR PAC-MAN is capable of inhibiting coronavirus fragment expression in human lung epithelial cells

To evaluate whether Cas13d is effective for targeting SARS-CoV-2 sequences, we created two reporters expressing synthesized fragments of SARS-CoV-2 fused to GFP (SARS-CoV-2-F1 and SARS-CoV-2-F2; **Fig. 2B, Supplemental Table 3**). As SARS-CoV-2 appears to mainly infect respiratory tract cells in patients^5^, we chose to use human lung epithelial A549 cells as a model cell line. We created a stable A549 cell line expressing Cas13d through lentiviral infection, followed by sorting for the mCherry marker that was co-expressed with Cas13d (**Fig. 2C**).

We first investigated which regions of the SARS-CoV-2 fragments were most susceptible to Cas13d-mediated targeting and cleavage. To do this, we transfected or transduced the SARS-CoV-2 reporters into Cas13d A549 cells that had been transduced with pools of four crRNAs targeting either RdRP or N gene regions (**Fig. 2D,** see **Methods**). We chose to evaluate the effectiveness of our system using a pool of crRNAs, since in the context of live viral infection, this would prevent the possibility of escape from inhibition due to a mutation involving a single crRNA target site. A total of ten groups (G1-G10) were tested for all 40 crRNAs (**Supplemental Table 2**). Twenty-four hours after SARS-CoV-2 reporter transfection, we performed flow cytometry to examine levels of GFP protein expression and collected RNA from cells to perform quantitative real-time PCR (qRT-PCR) to examine mRNA transcript abundance.

We observed that most RdRP-targeting crRNA pools were able to repress reporter expression to some extent compared to the control, and one pool targeting the middle of the SARS-CoV-2-F1 RdRP region (G4; SARS-CoV-2 genome coordinates 13,784-14,149) was able to repress GFP by 86% (p = 2×10^−6^) (**Fig. 3A, S2A-B, Supplemental Table 4**). One pool of N gene-targeting crRNAs (G8; SARS-CoV-2 genome coordinates 28,803-29,034) was able to substantially repress SARS-CoV-2-F2 GFP, causing a 71% decrease in expression (p = 2×10^−7^) (**Fig. 3B, S2A-B, Supplemental Table S4**). Our qRT-PCR analysis showed consistent results at the RNA level, with G4 and G8 inhibiting their respective reporter mRNA abundance by 83% (p = 3×10^−11^) and 79% (p = 2×10^−12^) (**Fig. 3A-B, Supplemental Table S4**). The variations in the ability of different crRNA pools to repress the SARS-CoV-2 reporters are likely due to RNA secondary structure inherent in the SARS-CoV-2 genome or differences in binding affinities for each crRNA sequence composition.

**Figure 3.**
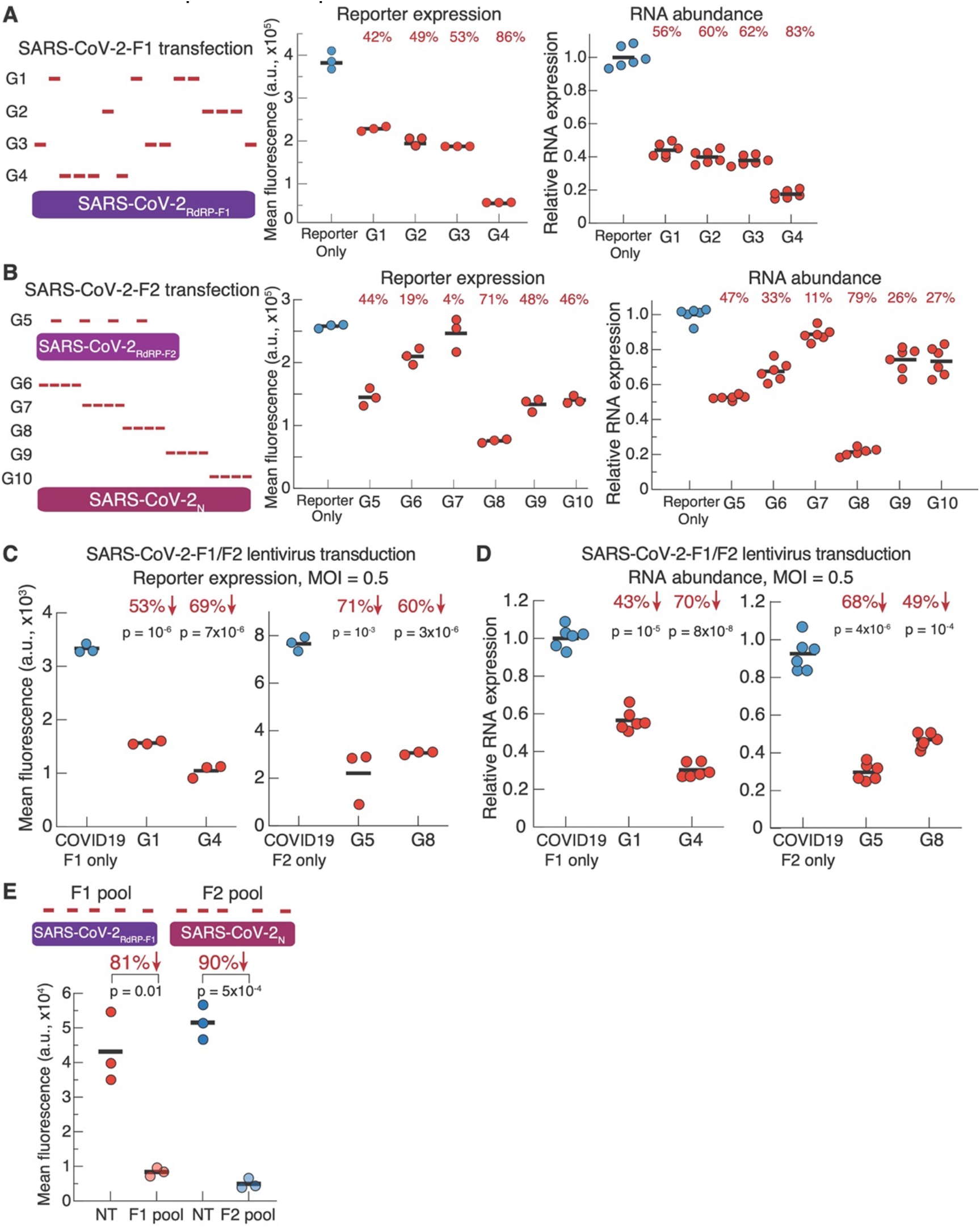
CRISPR PAC-MAN can effectively inhibit SARS-CoV-2 reporters. **(A-B)** Left, schematics of pools of crRNAs targeting the SARS-CoV-2-F 1 (**A**) or F2 (**B**) reporters for transfection. Middle, GFP expression as measured by flow cytometry. Right, mRNA abundance as measured by qRT-PCR. **(C)** GFP expression when SARS-CoV-2-F 1 (left) or SARS-CoV-2-F2 (right) is delivered via lentiviral transduction. MOI = 0.5. (**D**) RNA abundance when SARS-CoV-2-F1 (left) or SARS-CoV-F2 (right) is delivered via lentiviral transduction. MOI = 0.5. (**E**) GFP expression levels when using pools of crRNAs tiling across SARS-CoV-2-F1 or F2 reporters. Red, SARS-CoV-2-F1 reporter; blue, SARS-CoV-2-F2 reporter. NT, non-targeting crRNA controls. P values for **A-B** crRNA groups are shown in **Supplemental Table 4.** P values for **C-E** are labeled for each group.

We next independently validated the performance of adjacent crRNA pools when SARS-CoV-2-F1 or -F2 was introduced via lentiviral transduction. We argue that this approach, in which the lentivirus-derived RNA serves as a PAC-MAN target prior to lentivirus integration into DNA, mimics the crRNA targeting conditions that would apply for natural COVID-19 infection. We tested on a few selected crRNA pools, including the 3 best groups G4, G5, and G8 and one less efficient group G1, with a lentiviral multiplicity of infection (MOI) of 0.5. Our results revealed similar patterns to our SARS-CoV-2 transfection experiments, with the best groups (G4, G5, G8) able to repress their reporters by 69%, 71%, and 60% (p = 7×10^−6^, 10^−3^, 3×10^−6^ for G4, G5, G8, respectively) (**Fig. 3C**). qRT-PCR analysis also showed consistent results, with G4, G5, and G8 repressing 70%, 68%, and 49% of RNA abundance (p = 8×10^−8^, 4×10^−6^, 10^−4^ for G4, G5, G8 respectively) (**Fig. 3D**).

We further validated the effectiveness of Cas13d-mediated repression of SARS-CoV-2 reporters by co-transfecting each reporter with a pool of 5 crRNAs tiled along the whole fragment, compared to a pool of 5 non-targeting control crRNAs (**Table S2**). We found that the RdRP-targeting crRNAs were able to repress SARS-CoV-2-F1 GFP by 81% (p = 0.01), while N-targeting crRNAs were able to repress SARS-CoV-2-F2 GFP by 90% (p = 5×10^−4^) (**Fig. 3E, S3A**). We also did an analysis of the extent to which expression of Cas13d or crRNAs affected SARS-CoV-2 GFP reporter inhibition. Our analysis revealed that SARS-CoV-2 inhibition was more sensitive to crRNA concentration in our samples, while lower Cas13d expression only moderately decreased inhibitory activity (**Fig. S3B**). Thus, maintaining a high level of crRNA concentration is likely a determining factor for more effective SARS-CoV-2 sequence targeting and cleavage.

These data together suggest: 1) Cas13d PAC-MAN can be an effective system to target and degrade SARS-CoV-2 sequences in human cells and 2) proper design of crRNAs are important for obtaining a high efficiency of SARS-CoV-2 inhibition.

### CRISPR PAC-MAN is able to inhibit IAV infection in human lung epithelial cells

Since we do not yet have access to live SARS-CoV-2 virus, as a proof-of-concept we elected to test the CRISPR-Cas13d strategy on inhibiting H1N1 IAV, a RNA virus with a similar tropism as SARS-CoV-2 for respiratory tract epithelial cells. In contrast to SARS-CoV-2, which has only a single, continuous RNA genome, the IAV genome is contained in 8 negative-sense RNA segments, in which each plays an important role in packaging the viral RNA into budding virions^16,17^.

To create crRNAs targeting a broad range of IAV strains, we aligned all complete IAV genomes retrieved from the Influenza Research Database and searched for the most conserved regions (**Supplemental Table 5**). Our analysis and the work of others showed that the segment ends are highly conserved for all 8 segments^16-21^. The segment ends are potentially attractive regions for targeting with Cas13d because of the potential to inhibit a broad range of IAV strains and because of the previous evidence that interfering with the packaging of one segment encoded on the segment ends can decrease the packaging efficiency of other segments and overall virion packaging^18,22-25^. Using sequence representation from ~140 different strains of influenza, we designed crRNAs that could be robust across as many different IAV strains as possible. Our bioinformatics analysis resulted in 6 crRNAs targeting highly conserved genome regions for each of the 8 IAV segments, for a total of 48 crRNAs (**Supplemental Table 6**).

To test the efficiency of IAV-targeting crRNAs, we performed a screen of our crRNA pools in the stable Cas13d A549 cell line to determine which were most effective at inhibiting IAV infection (**Fig. 4A**). To do this, we transfected a pool of 6 crRNAs targeting a given IAV genome segment. Two days after crRNA transfection, the Cas13d A549 cells were challenged with PR8-mNeon, a strain of H1N1 IAV (A/Puerto Rico/8/1934) engineered to express the mNeonGreen gene, a fluorescent reporter protein (hereafter referred to as PR8-mNeon) at an MOI of 2.5 or 5.0^26-28^. At approximately 18 hours post-challenge, the cells were analyzed for IAV infection through flow cytometry and microscopy. We examined the percentage of cells that were mNeon positive for each segment-targeting crRNA pool and calculated the fold-change compared to cells transfected with a pool of non-targeting crRNAs).

**Figure 4.**
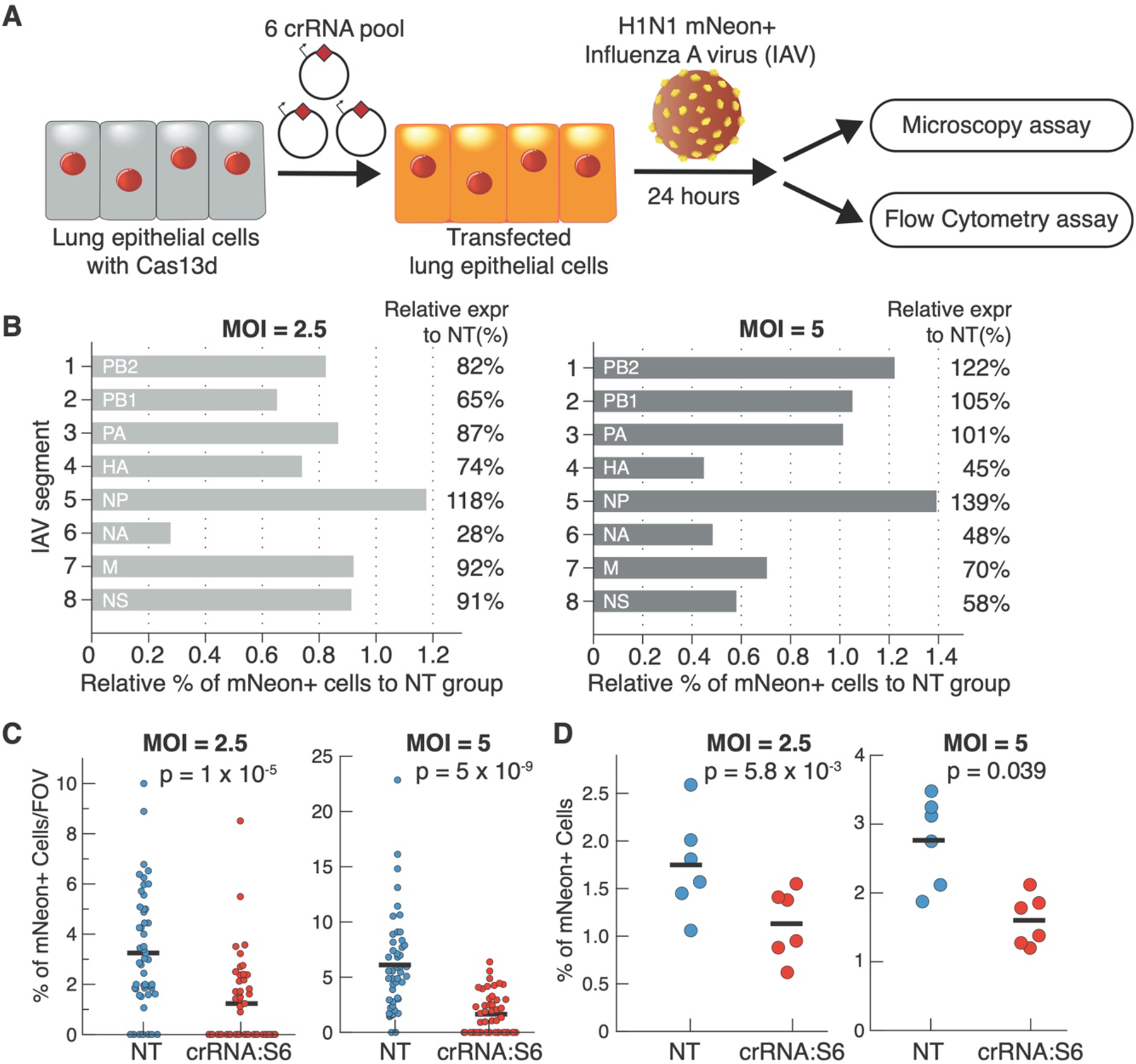
Inhibition of IAV infection using CRISPR PAC-MAN. **(A)** Workflow used to challenge Cas13d A549 lung epithelial cells with PR8 mNeon IAV. **(B)** Screen of pools of 6 crRNAs targeting each of the eight IAV genome segments. % of mNeon+ cells in each of the eight crRNA conditions are compared to a pool of non-targeting (NT) crRNAs at an MOI of 2.5 (left) or 5 (right). **(C)** Microscopy quantification of % of mNeon+ cells per field of view (FOV) at an MOI of 2.5 (p = 1×10^−5^, left) and 5 (p = 5×10^−9^, right). Each dot represents the % for a single microscopy FOV. N = 48 FOV. **(D)** Flow cytometry evaluation of % of mNeon+ cells at an MOI of 2.5 (p = 0.04, left) and 5 (p = 0.006, right). N = 6 biological samples.

Out of all crRNAs, the crRNA pool targeting segment 6 (S6) delivered the most consistent and robust results across different MOIs (72% reduction for MOI = 2.5 and 52% reduction for MOI = 5; **Fig. 4B, S4**). Additionally, the segment 4 (S4) pool also showed moderate but consistent inhibition. To verify Cas13d-mediated inhibitory effects on S4 and S6, we further tested at a lower MOI of 0.5, which showed better inhibition of IAV across samples (e.g., 79% reduction using crRNAs targeting S6; **Fig. S5**). This confirms consistent inhibitory activity of IAV by Cas13d when targeting these two segments. There is a possible trend of greater inhibition at lower viral titers, suggesting that Cas13d PAC-MAN could be a particularly effective strategy for prevention of new infections, which typically occurs from exposure to a low level of virus.

We further characterized the S6-targeting crRNA pool. S6 encodes the gene neuraminidase (NA), which releases budding virions from the host cell ^16^. We performed microscopy quantification of mNeon+ cells and found that mNeon expression was reduced by 62% (p = 10^−5^) and 73% (p = 5×10^−9^) with an MOI of 2.5 and 5, respectively, with S6-targeting crRNAs (**Fig. 4C**) and this trend was corroborated by flow cytometry (**Fig. 4D**). Together, our data show that Cas13d PAC-MAN is able to target highly conserved viral regions and robustly inhibit viral replication in human lung epithelial cells.

### Cas13d PAC-MAN as a potential pan-coronavirus inhibition strategy

In the past two decades, multiple variants of coronavirus, including those causing COVID-19, SARS and MERS, emerged from animal reservoirs and infected humans, each time causing significant disease and fatalities^1-3^. Thus, the design of a strategy that could broadly prevent and target viral threats, including all coronavirus strains that currently reside in animals, would be a valuable resource. We asked whether it is possible to design a minimal number of crRNAs that could target a majority of known coronaviruses found in both humans and animals.

To do this, we analyzed all known coronavirus genomes (3,051) to identify all possible crRNAs that can target each of those genomes, ending up with approximately 6.48 million possible crRNAs (**Fig. 5A, S6**). We then refined our list to define the smallest number of crRNAs that could target all known coronavirus genomes with zero mismatches. We found that just two crRNAs could target ~50% of coronavirus genomes, including those causing COVID-19, SARS, and MERS; 6 crRNAs were able to target ~91% of coronavirus genomes; and 22 crRNAs covered all sequenced coronaviruses with no mismatches (**Fig. 5B, Supplemental Table 7**). The ability to use a relatively small number of crRNAs to broadly target most or all coronavirus strains further highlights the uniqueness of our approach in contrast to traditional pharmaceutical or vaccination approaches.

As a resource, we provide a map to show how our top 6 crRNAs (PAC-MAN-T6) targeted the coronavirus phylogenetic tree (**Fig. 5A**, red box). Beta coronaviruses, the genus of viruses causing COVID-19, SARS, and MERS, were mostly covered by 3 crRNAs (crRNA 1-3), while the other genera were covered by these three plus the remaining 3 crRNAs. Altogether, the PAC-MAN-T6 can target all known human coronavirus with broad coverage against other animal coronaviruses.

The minimal set of six crRNAs also included one crRNA (designated crRNA-N18f) that we validated. A recent work analyzed 103 SARS-CoV-2 genomes and indicated that there are two major types (L and S) of SARS-CoV-2 existing with different viral signature sequences^4^. To verify whether crRNA-N18f can target all L and S strains, we retrieved a set of 202 recently sequenced SARS-CoV-2 sequences from Global Initiative on Sharing All Influenza Data (GISAID). Notably, crRNA-N18f targets all 201 genomes, with only one SARS-CoV-2 sequence carrying a single mismatch mutation (**Supplemental Table 8**). This suggests that our approach can robustly target multiple SARS-CoV-2 strains, but also suggests the potential need to use a pool of crRNAs to avoid mutational escapes.

**Figure 5.**
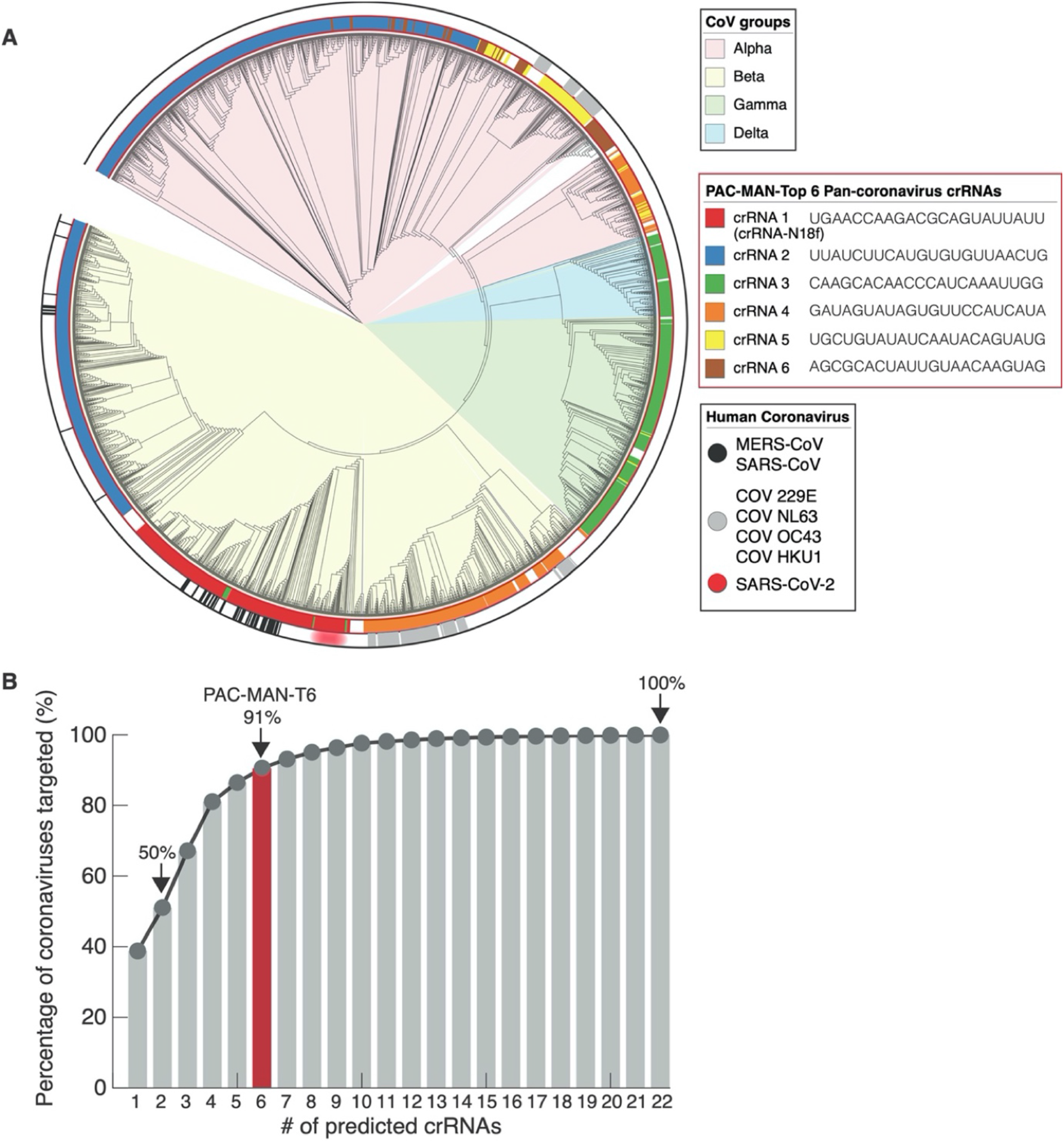
Pan-coronavirus targeting using a minimal pool of PAC-MAN crRNAs. **(A)** A view of the phylogenetic tree of all sequenced coronaviruses, organized by genus of strains. The inner ring depicts coverage by each of the top 6 (PAC-MAN-T6) crRNAs targeting all human coronaviruses. The outer ring shows current species of coronaviruses that are infectious to humans, including COVID-19 (red). **(B)** A histogram showing the minimum number of predicted crRNAs it takes to cover a maximum number of coronavirus genomes.

## DISCUSSION AND PERSPECTIVES

CRISPR-Cas13 emerged as a natural bacterial defense against bacteriophages, a powerful antiviral system that is able to use sequence-specific crRNAs to protect host bacterial cells from infections^29^. Here, we have repurposed the RNA-guided RNA endonuclease activity of Cas13d in mammalian cells against emergent viral targets, SARS-CoV-2 and IAV, in our PAC-MAN strategy. This expands the applications of CRISPR-Cas13 systems in addition to their uses for diagnostics such as SHERLOCK and live-cell imaging^30-33^. Prior work showed Cas13a/b systems could inhibit ssRNA viruses including influenza, which is consistent with our observed high efficiency of SARS-CoV-2 reporter and IAV inhibition using the PAC-MAN approach^31^. We demonstrated that Cas13d-based genetic targeting can effectively target and cleave the RNA sequences of SARS-CoV-2 fragments and IAV in lung epithelial cell cultures. Our bioinformatics analysis also suggested a minimal pool of 6 crRNAs able to target over 90% of sequenced coronaviruses. While this strategy is a proof-of-concept and will require further testing using replication-competent SARS-CoV-2 viruses and validation in animal models before clinical tests in humans, it represents a unique approach to implement a rapid and broad antiviral defense in humans against emerging pathogens for which there are no effective vaccines.

For both SARS-CoV-2 and IAV, we found highly conserved regions of the viral genomes to target with Cas13d. In the case of SARS-CoV-2, these genome regions encode the RdRP and Nucleocapsid proteins, which are essential for coronavirus replication and function^13,34^. RdRP is responsible for catalyzing the replication of all viral mRNAs and Nucleocapsid binds to genomic RNA, protecting it and serving as one of the two main structural proteins in virions. Targeted inhibition of these proteins could have an outsized effect on disabling virus production and function, in addition to reducing viral load through degradation of the viral genome itself. For IAV, the crRNAs that showed the best viral inhibition target the conserved ends of the segment S6 that encodes Neuraminidase (NA), a viral surface protein essential for mediating budding of new virions^35^. Prior mutation analysis has shown that mutating the packaging signals of one segment end leads to decreased packaging of other segments, indicating that there is a cooperative mechanism for packaging the IAV genome^18^. Thus, by inhibiting a single IAV genome segment, we could reduce the overall packaging of IAV in a synergistic fashion.

Notably, the SARS-CoV-2 RdRP gene and IAV NA gene that we targeted in our study are also the targets of two antiviral drugs, Remdesivir and Oseltamivir. Remdesivir, created by Gilead Sciences (development code GS-5734), is an RdRP inhibitor that was developed to treat Ebola and Marburg virus infections and has been reported to have antiviral activity against MERS and SARS^36-38^. Remdesivir is already administered for COVID-19 through compassionate use requests, and clinical trials for its use against COVID-19 have already started in the U.S. (clinical trial ID NCT04280705) and China. Oseltamivir (also known as Tamiflu), is an antiviral medication for treating and preventing IAV that acts as an inhibitor of NA, thereby preventing the egress of new viral particles^39^. It will be interesting to see if these drugs could be used in combination with Cas13d PAC-MAN to synergistically cripple viral function and replication.

While we demonstrate that the PAC-MAN system is able to exhibit strong repression of viral sequences in an in vitro setting, it must be coupled with an effective in vivo delivery system for therapeutic use. There are several potential strategies that could be employed for transient in vivo expression of CRISPR components. Cas13d and its cognate crRNAs could be delivered as RNAs within polymers or lipid nanoparticles, perhaps with chemical alterations to increase stability^40^. In addition, a DNA-based liposomal delivery strategy, such as the recently developed HEDGES platform is also attractive^41^. Yet another strategy would be to deliver a ribonucleoprotein complex containing purified Cas13d assembled with crRNAs^42,43^. All of these could potentially be used both as a therapeutic and as a prophylactic measure.

If the delivery barriers are overcome and a strategy like this is able to be used therapeutically, there are many potential benefits over traditional vaccines. Traditional vaccines rely on priming the immune system through exposure to viral proteins or peptides often derived from surface proteins that exhibit a high rate of mutation, which increased the chances of viral evasion of the host immune response^44^. In contrast, here we have demonstrated a genetic strategy that is able to target highly conserved regions, which would be expected to make it much more unlikely for the virus to escape inhibition through mutation. In addition, the ability of Cas13d to process its own crRNAs from a crRNA array means that multiple crRNAs targeting different regions (e.g. **Fig. 3E,** and PAC-MAN-T6 in **Fig. 5A**) could be delivered simultaneously^9^, further reducing the chances of viral escape. As a further advantage, we demonstrate a potential pan-coronavirus strategy to target not only viruses that circulate in humans, but also those that currently are found in animal reservoirs that might transfer to humans to cause disease. If the crRNAs targeting these viruses can be tested and validated before they ever infect humans, we can greatly accelerate the development of countermeasures for future emergent threats.

In summary, our PAC-MAN strategy represents a potentially powerful new approach for inhibiting viral function and replication, and we envision it could be useful for a diverse array of circulating and emergent viral threats.

## ACKNOWLEDGEMENTS

The authors thank all members from Lei S. Qi lab and David Lewis lab for facilitating experiments and useful discussions. The authors thank the researchers who generated and contributed the SARS-CoV-2 sequence data to GISAID (https://www.gisaid.org/). The project is supported by a contract grant from Defense Advanced Research Projects Agency (DARPA) (Grant # HR001119C0060), as well as a Li Ka Shing Foundation gift fund (to Lei S. Qi lab).

## AUTHOR CONTRIBUTIONS

T.R.A., M.L., D.B.L., and L.S.Q. conceived of the idea. T.R.A., G.D., Y.L., L.G., L.Z., M.L., D.B.L., and L.S.Q. planned the experiments. T.R.A., X.L., and L.G. designed crRNAs. T.R.A., L.G., and T.P. cloned crRNAs. Y.L. designed and cloned SARS-CoV-2 reporters. T.R.A., Y.L., L.G., and L.Z. performed SARS-CoV-2 reporter cell culture experiments. T.R.A., G.D. performed IAV cell culture experiments. S.C., N.H., R.D. and D.R. provided key suggestions or facilitated IAV viral challenge experiments. X.L. and A.C. performed SARS-CoV-2 bioinformatics analyses. T.R.A. performed IAV bioinformatics analyses. T.R.A., G.D., Y.L., L.G., M.L., D.L. and L.S.Q. analyzed the experimental data. M.L., D.B.L. and L.S.Q. wrote the manuscript. All authors read and commented on the manuscript.

## COMPETING FINANCIAL INTERESTS

There are provisional patents filed via Stanford University related to the work.

## COMPUTATIONAL AND EXPERIMENTAL METHODS

### Genome Sequences Collection

47 complete SARS-CoV-2 assemblies were downloaded from (https://www.ncbi.nlm.nih.gov/genbank/2019-ncov-seqs/) on February 21, 2020. All the other 3,108 Coronavirinae complete genome sequences were downloaded from ViPR (Virus Pathogen Resource) database on February 21, 2020. After excluding very short genomes, we collected in total 3,051 Coronavirinae virus genomes. 202 SARS-CoV-2 genome sequences were downloaded from GISAID (Global Initiative on Sharing All Influenza Data; https://www.gisaid.org/).

### High-throughput crRNA Design

We developed a full computational workflow to design crRNAs for the Cas13d system. To design all possible crRNAs for the three pathogenic RNA viruses (SARS-CoV-2, SARS-CoV, and MERS-CoV), the reference genomes of SARS-CoV, MERS-CoV, along with SARS-CoV-2 genomes derived from 47 patients were first aligned by MAFFT using the --auto flag. crRNA candidates were identified by using a sliding window to extract all 22 nt sequences with perfect identity among the SARS-CoV-2 genomes. We annotated each crRNA candidate with the number of mismatches relative to the SARS-CoV and MERS-CoV genomes, as well as the GC content. 3,802 crRNA candidates were selected with perfect match against the 47 SARS-CoV-2 genomes and with ≤1 mismatch to SARS-CoV or MERS-CoV sequences. To characterize the specificity of 22nt crRNAs, we ensured that crRNA do not target any sequences in the human transcriptome. We used Bowtie 1.2.2 to align crRNAs to the human transcriptome (HG38; including non-coding RNA) and removed crRNAs that mapped to the human transcriptome with fewer than 2 mismatches.

### Computational identification of minimal set of crRNAs for pan-coronavirus targeting

We maximized the vertical coverage of pooled crRNAs to target a maximal set of the 3,051 Coronavirinae viruses with a minimal set of crRNAs. A sliding window was used to extract all unique 22 nt sequences in the Coronavirinae genomes, yielding 6,480,955 unique crRNA candidates. A hash table was used to map each crRNA candidate to the number of Coronavirinae genomes that it targets with a perfect match. A greedy algorithm was used to identify a minimal set of covering crRNAs. In each iteration, the crRNAs were sorted in descending order by the number of genomes targeted. The crRNA that targeted the most genomes was added to the minimal set of crRNAs, and all of the genomes it targeted were removed from the hash table values. If multiple crRNAs were tied for targeting the most genomes, the crRNAs were ranked by the length and quantity of stretches of repeated nucleotides, and the lowest ranking crRNA (i.e. the one with the fewest, shortest stretches of repeated nucleotides) was added to the set of minimal crRNAs. Using this approach, a set of 22 crRNAs was identified that was able to target all 3150 Coronavirinae virus genomes (including 47 SARS-CoV-2). The phylogenetic tree and the associated annotations were generated by the Interactive Tree Of Life (iTOL v5.5; https://itol.embl.de/).

### Conservation Plot Creation

All the complete SARS-CoV-2 genome sequences were aligned with the SARS-CoV and MERS-CoV NCBI Refseq sequences by MAFFT using the --auto flag. The alignment was visualized using the Geneious Prime software (Geneious 2020.0.4; https://www.geneious.com).

A consensus nucleotide (which is shared by >50% of non-gap characters at the given position) was assigned to each position in the alignment. Not every position has a consensus nucleotide. The percent conservation at each position was calculated as the percentage of sequences matching the consensus at that position. To aid interpretation, the resulting plot of percentage conservation was smoothed using a moving average with a window size of 1,001 bp centered at each position. Positions corresponding to gap characters in the SARS-CoV-2 reference sequence were omitted.

### Design and cloning of RdRP- and Nucleocapsid-targeting crRNAs

The crRNA plasmids were cloned using standard restriction-ligation cloning. Forward and reverse oligos for each spacer (IDT) were annealed and inserted into backbone using T4 DNA Ligase (NEB). Assembled oligos were either inserted into a pHR or pUC19 backbone, obtained and modified from Addgene (#121514 and #109054, respectively). Spacer sequences for all crRNAs can be found in **Supplemental Table 2 and 6.**

### Design and cloning of SARS-CoV-2 reporters

To clone SARS-CoV-2 reporters, pHR-PGK-scFv GCN4-sfGFP (pSLQ1711) was digested with EcoR1 and Sbf1 and gel purified. The PGK promoter and sfGFP were PCR amplified from pSLQ1711 and gel purified. The SARS-CoV-2-F1 and SARS-CoV-2-F2 fragments were synthesize from Integrated DNA Technologies (IDT). At last, the PGK, sfGFP, and SARS-CoV-2-F1/F2 fragments were inserted into the linearized pSLQ1711 vector using In-Fusion cloning (Takara Bio).

### Lentiviral packaging and stable cell line generation

On day 1, one confluent 100mm dish (Corning) of HEK293T cells were seeded into three 150mm dishes (Corning). On day 2, cells were ~50-70% confluent at the time of transfection. For each dish, 27.18 μg of pHR vector containing the construct of interest, 23.76 μg of dR8.91 and 2.97 μg of pMD2.G (Addgene) were mixed in 4.5 mL of Opti-MEM reduced serum media (Gibco) with 150 μL of Mirus TransIT-LT1 reagent and incubated at room temperature for 30 minutes. The transfection complex solution was distributed evenly to HEK293T cultures dropwise. On day 5, lentiviruses are harvested from the supernatant with a sterile syringe and filtered through a 0.22-μm polyvinylidene fluoride filter (Millipore).

For A549 cells, lentivirus precipitation solution (Alstem) was added and precipitated as per the manufacturer’s protocol. One well of a 6-well plate (Corning) of A549 cells at ~50% confluency were transduced with the precipitated lentivirus. After 2-3 days of growth, the cell supernatant containing virus was removed and the cells were expanded. Cells were then sorted for mCherry+ cells using a Sony SH800S cell sorter.

### SARS-CoV-2 reporter challenge experiments

Fig. 3A-3D: On day 1, A549 cells stably expressing were seeded at a density of 60,000 cells per well in a 12-well plate. On day 2, cells were transduced with pooled crRNAs targeting specific regions on SARS-CoV-2 (Supplemental Table S2). On day 3, cells were switched to fresh medium with Puromycin at 2 ug/mL. Twenty-four hours after Puromycin selection, cells with transfected crRNAs were challenged with SARS-CoV-2 reporter plasmids or SARS-CoV-2 reporter lentivirus. Twenty-four hours after transfection or 48 hours after transduction, cells were tested using flow cytometry on a Beckman-Coulter CytoFLEX S and qRT-PCR on a Biorad

CFX384 real-time system. Statistical analyses were done using a two-sided t-test with unequal variance in Excel to calculate p values.

**Fig. 3E:** On day 0, 30,000 wildtype A549 cells and 30,000 A549 cells stably expressing Cas13d (hereafter referred to as Cas13d A549) were plated in 24-well plates (Corning). On day 1, Cas13d A549 cells were transfected with two pools of five crRNAs targeting the two SARS-CoV-2 reporters or a non-targeting crRNA along with the respective SARS-CoV-2 reporters in an equimolar ratio using Mirus LT1 reagent (50 μL Opti-MEM reduced serum media, 0.136 μg crRNA pool, 0.364 μg SARS-CoV-2 reporter, 1 μL Mirus LT1 per well). Wildtype cells were transfected with either the crRNA pools or SARS-CoV-2 reporter plasmids as compensation controls. GFP expression was measured on day 3 by flow cytometry on a Beckman-Coulter CytoFLEX S instrument. Statistical analyses were done using a two-sided t-test with unequal variance in Excel to calculate p values.

### Transduction of SARS-CoV-2 reporter plasmid

On day 0, Lenti-X 293T cells (Takara Bio) were seeded to 1:6 from a 100 mm dish in a 150 mm dish. The following day, Lenti-X 293T cells were transfected with 27.18 μg of SARS-CoV-2 reporter, 23.76 μg of dR8.91 and 2.97 μg of pMD2.G (Addgene) in 4.5 mL of Opti-MEM reduced serum media (Gibco) with 150 μL of Mirus TransIT-LT 1 reagent. 30,000 Cas13d A549 and wildtype A549 cells were also seeded in 24-well plates. On day 2, A549 cells were transfected with pools of targeting crRNAs or a non-targeting guide (50 μL Opti-MEM reduced serum media, 0. 5 μg crRNA pool, 1 μL Mirus LT1 per well). On day 3, lentiviruses are harvested from the supernatant with a sterile syringe and filtered through a 0.22-μm polyvinylidene fluoride filter and precipitated with lentivirus precipitation solution and resuspended in 7 mL DMEM media. The media on the cells was replaced with either fresh media for the compensation controls or 0.5 mL virus-containing media at the following dilutions for the experimental conditions: undiluted, 1/2 dilution, 1/3 dilution, and 1/4 dilution. On day 4, the media on all the cells was replaced with 0.5 mL fresh DMEM. On day 5, cells were measured by flow cytometry and assessed for GFP knockdown. Statistical analyses were done using a two-sided t-test with unequal variance in Excel to calculate p values.

### Generation and titering of PR8 mNeon IAV

The PR8 mNeon IAV was generated by insertion of the mNeon fluorescent gene into segment 4 of the H1N1 A/Puerto Rico/8/1934 strain (Nicholas Heaton’s Lab, Duke University) as previously described (Heaton et al., 2017). The PR8-mNeon virus was propagated in chicken eggs, and harvested allantoic fluid containing the virus was titered using MDCK plaque assays, and frozen in aliquots for later use.

### IAV challenge experiments

3.1×10^6^ Cas13d A549 cells were plated in two 100 mm dishes. The following day, the cells were transfected with 15 μg pooled crRNAs targeting the IAV genome or non-targeting controls (Supplemental Table 6) in 1.5 mL Opti-MEM reduced serum media and 30 μL Mirus LT1 reagent. After approximately 24 hours, the transfected Cas13d A549 cells were dissociated and re-plated in 24-well plates at a density of 15-20,000 cells/well and incubated overnight. Flow cytometry was performed on excess cells to quantify the transfection efficiency. Cells were challenged overnight with mNeonGreen expressing H1N1 A/Puerto Rico/8/1934 at an MOI of 2.5 and 5. Uninfected cells were included as a negative control. An independent two-tailed T test was used to calculate the p values and statistics in **Fig. 4C-D**.

### Quantitative RT-PCR (qRT-PCR)

Real-time qRT-PCR was performed to quantify RNA abundance. For each sample, total RNA was isolated by using the RNeasy Plus Mini Kit (Qiagen Cat# 74134), followed by cDNA synthesis using the iScript cDNA Synthesis Kit (BioRad, Cat# 1708890). qRT-PCR primers were ordered from Integrated DNA Technologies (IDT). Quantitative PCR was performed using the PrimePCR assay with the SYBR Green Master Mix (BioRad), and run on a Biorad CFX384 realtime system (C1000 Touch Thermal Cycler), according to manufacturers’ instructions. Cq values were used to quantify RNA abundance. The relative abundance of the SARS-CoV-2 fragments was normalized to a GAPDH internal control.

### Microscopy

Microscopy for SARS-CoV-2 experiments was performed on a Nikon TiE inverted confocal microscope equipped with an Andor iXon Ultra-897 EM-CCD camera and 405 nm, 488 nm, 561 nm and 642 nm lasers, using the 20x PLAN APO air objective (NA = 0.75). Image were taken using NIS Elements version 4.60 software. Microscopy for IAV experiments was performed on a Keyence BZ-X810 microscope equipped with a GFP and Texas Red filter was used to obtain images with a 20X objective lens.

### Flow Cytometry

A549 cells that express Cas13d and are transfected with crRNAs and challenged with PR8 mNeon virus were washed with 1X PBS, treated with 1X Trypsin EDTA and fixed in 1.6% paraformaldehyde in PBS for 10 minutes. Cells were washed with FACS buffer (1X PBS, 0.3% BSA, 1mM EDTA). BD Accuri C6 plus flow cytometer was used to collect raw data. Raw data were processed with Cytobank (BD) software.

