## Supplemental Figures and Information for "Development of CRISPR as a prophylactic strategy to combat novel coronavirus and influenza"

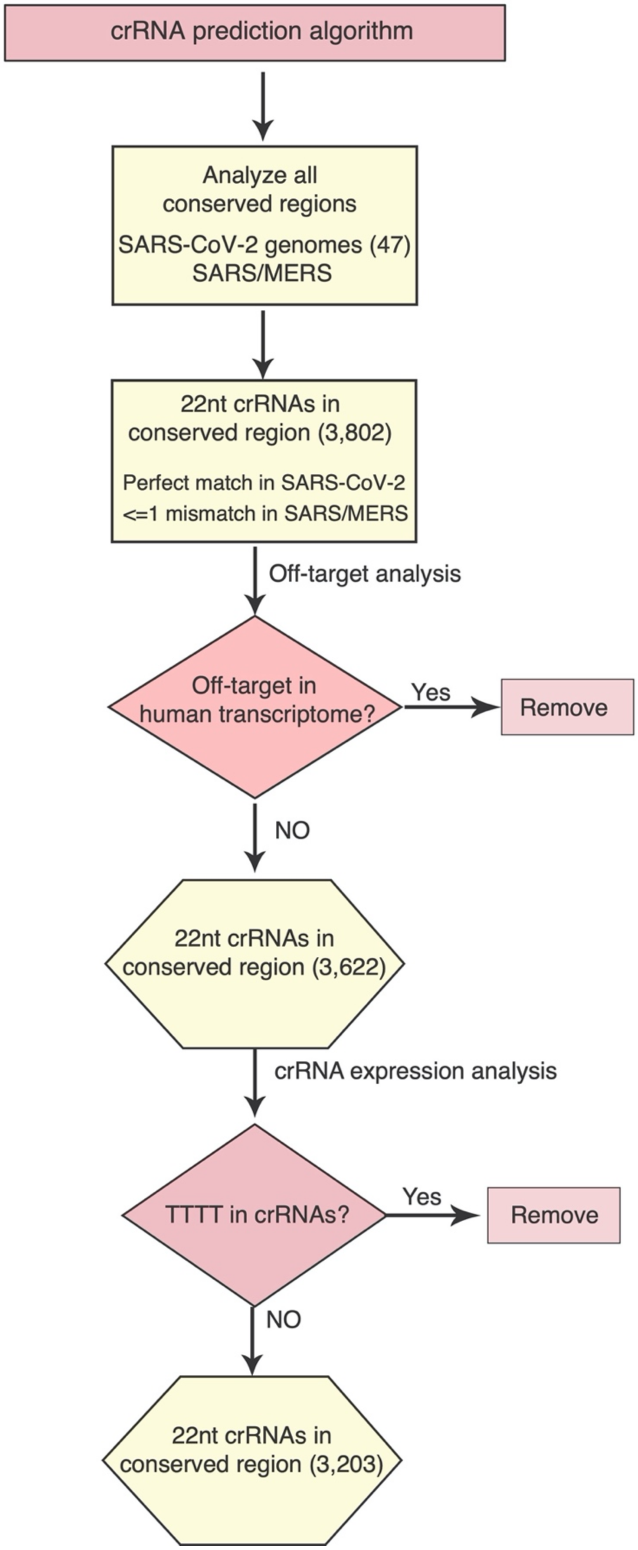


**Figure S1 | A bioinformatic pipeline to predict effective and specific crRNA designs.** Our bioinformatic method analyzes all possible crRNAs that target regions conserved between reported SARS-CoV-2, SARS-CoV, and MERS-CoV.

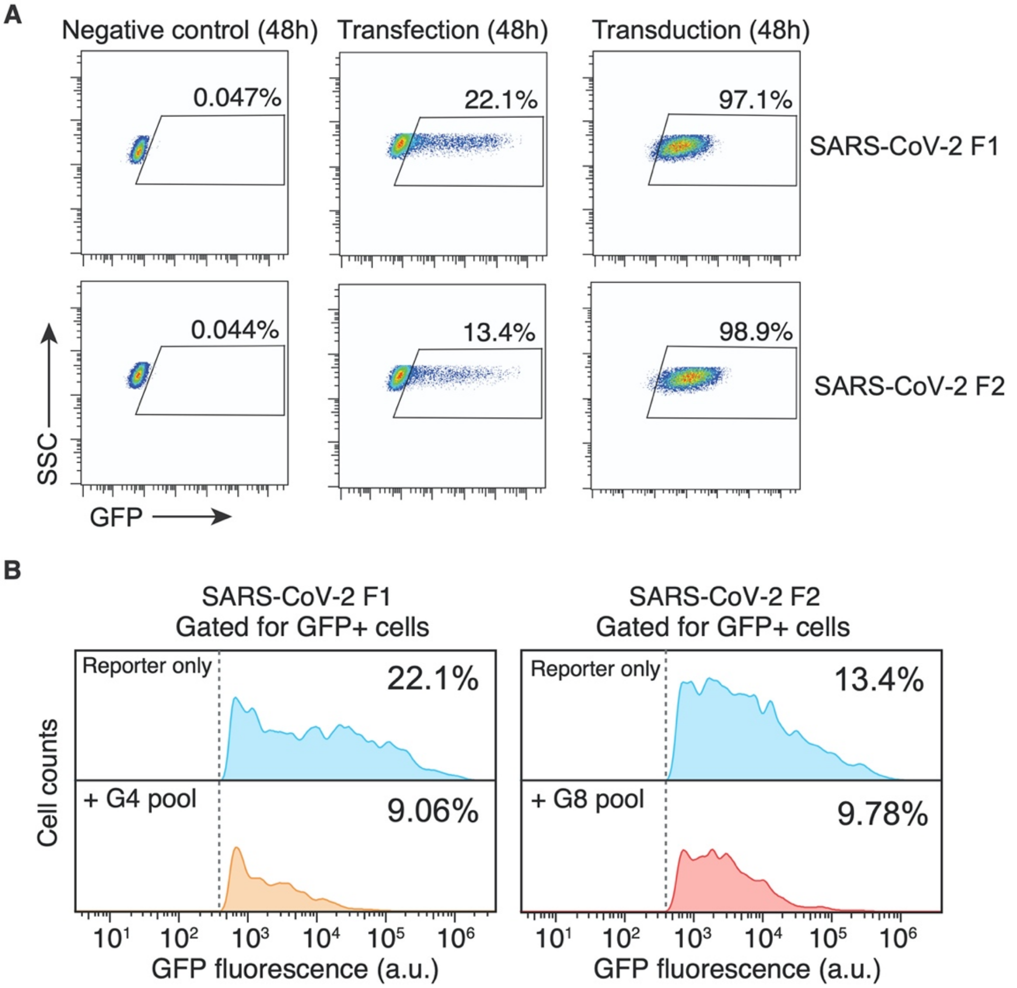


**Figure S2 | Flow cytometry plots demonstrating SARS-CoV-2-19 F1 and F2 transfection and transduction. (A)** Examination of SARS-CoV-2-19 reporter levels in A549 cells 48 hours after transfection or transduction. While transfection led to a much lower percentage of GFP positive cells compared to transduction, transfection also led to much higher levels of GFP expression, giving a better dynamic range of repression. **(B)** Representative flow histograms of GFP reporter levels after SARS-CoV-2 reporter challenge. Cells shown were gated for GFP+ cells only.

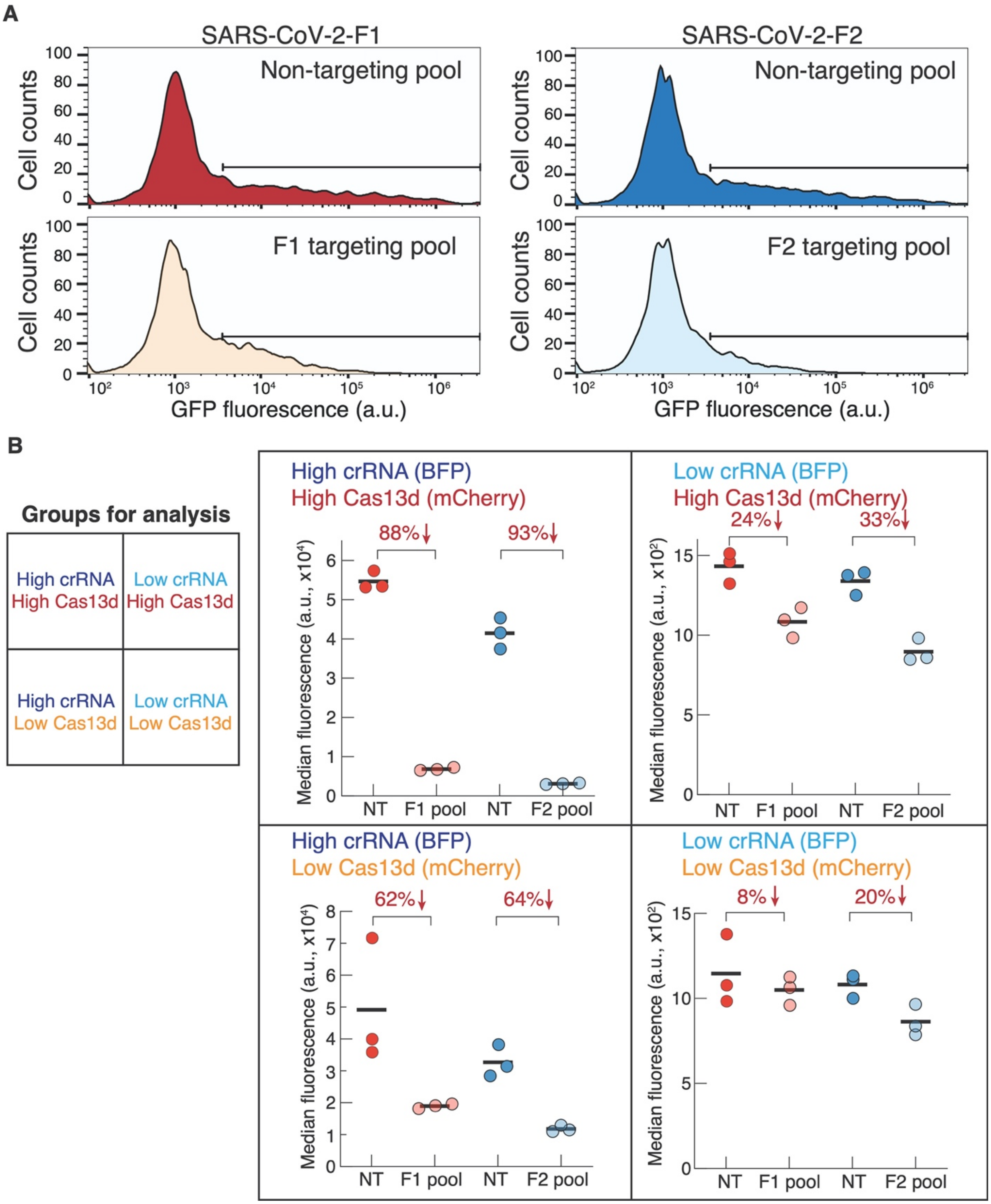


**Figure S3 | Flow cytometry plots demonstrating tiled crRNAs with a strong repression of the SARS-CoV-2 F1 and F2 reporters.** (**A**) Flow cytometry histograms of GFP expression after SARS-CoV-2-F1 (left) and SARS-CoV-2-F2 (right) reporter challenge. (**B**) Analysis of different expression levels of Cas13d (marker: mCherry) and crRNA pools (marker: BFP) and the resulting effects on SARS-CoV-2-F1/F2 reporter inhibition. Percent reduction of GFP expression is labeled on the top. Red, SARS-CoV-2-F1 reporter; blue, SARS-CoV-2-F2 reporter.

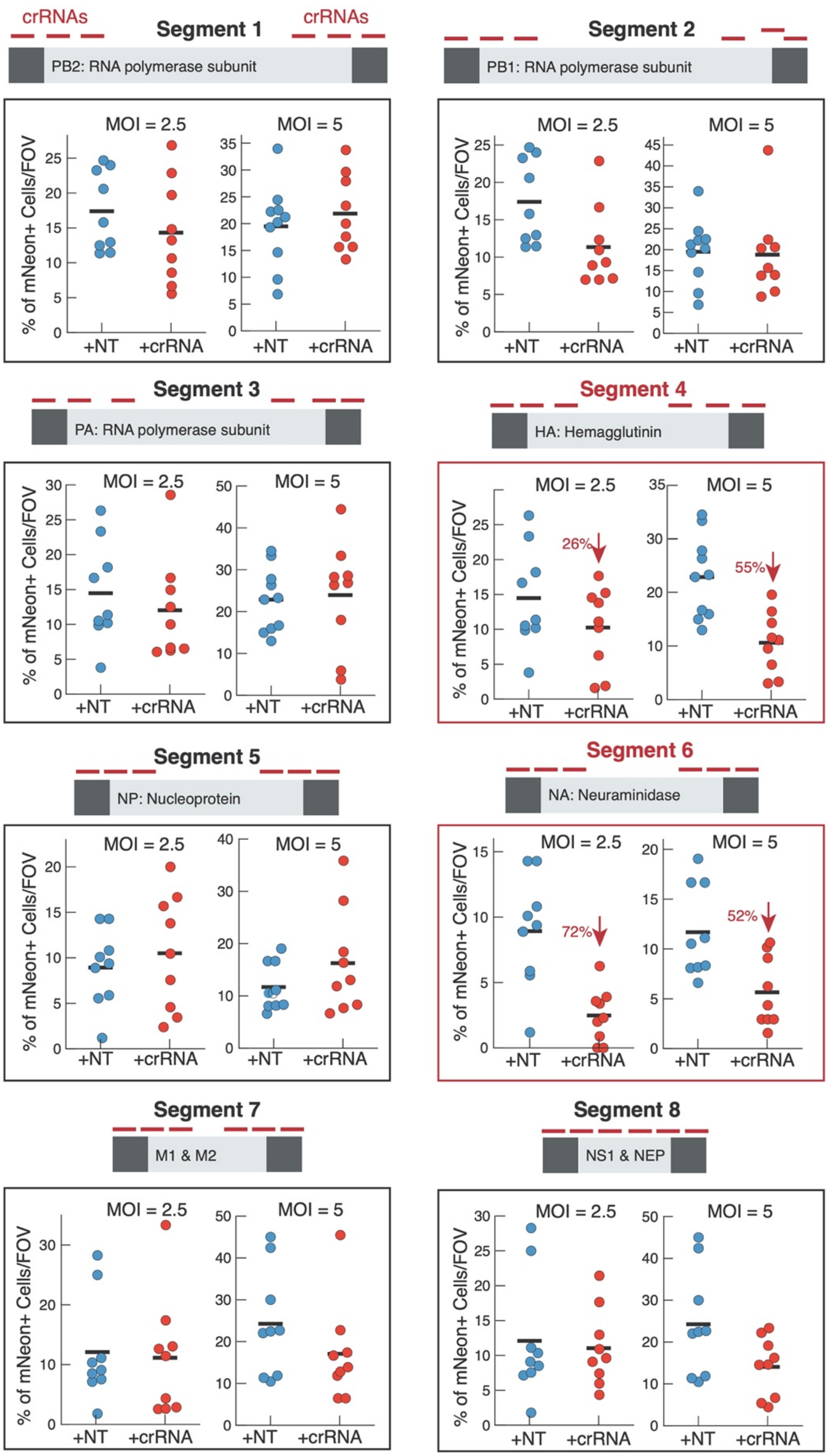


**Figure S4 | Screening of pools of 6 crRNAs targeting each of the eight IAV genome segments.** Each panel shows the percentage of quantified mNeon+ cells under two infection conditions (MOI = 2.5 or 5). Each dot represents the percentage for a single microscopy field of view (FOV). Blue, non-targeting (NT) crRNAs; red, targeting crRNAs.

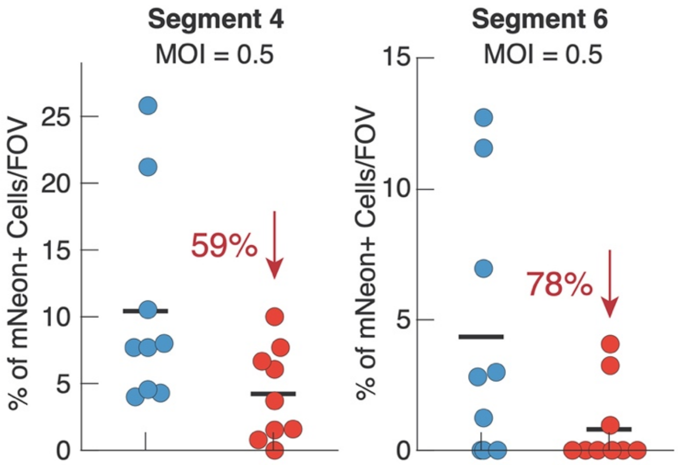


**Figure S5 | Testing of pools of 6 crRNAs targeting IAV S4 and S6 using a lower MOI of 0.5.** Each dot represents the percentage for a single microscopy field of view (FOV). Blue, non-targeting (NT) crRNAs; red, targeting crRNAs.

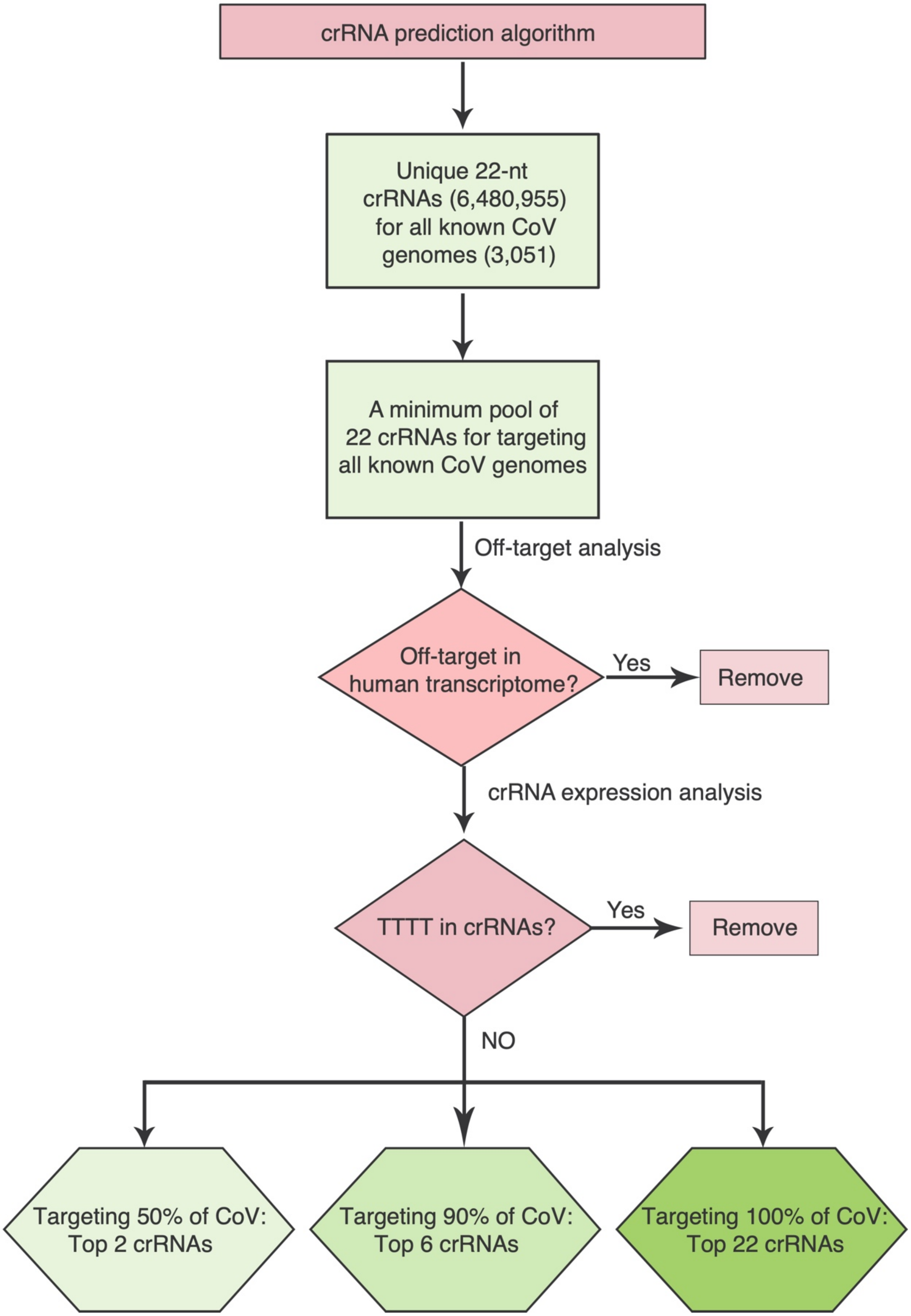


**Figure S6 | A bioinformatic pipeline to predict minimum sets of pan-coronavirus crRNA designs.** The bioinformatic pipeline analyzed and predicted pan-coronavirus targeting crRNAs. The number of crRNAs at each step of the workflow is denoted in parentheses.

### SUPPLEMENTAL TABLES

**Supplemental Table 1 | Sequences of 3,203 predicted crRNAs targeting SARS-CoV-2 and their coordinates**

**Supplemental Table 2 | Sequences of 40 crRNAs targeting SARS-CoV-2 and non-targeting crRNAs**

**Supplemental Table 3 | Synthesized SARS-CoV-2 fragment sequences**

**Supplemental Table 4 | P values for screening pools of 40 crRNAs in Figure 3A-B**

**Supplemental Table 5 | Weblogos of IAV Segments**. All complete, pre-aligned, complementary RNA (cRNA) segment sequences for IAV were downloaded from the Influenza Research database. All sequences were reverse-transcribed to viral RNA (vRNA) and filtered for the most-frequently occurring length for each segment. Each filtered segment was uploaded to Weblogo3 (http://weblogo.threeplusone.com/) to create weblogos for visualization.

**Supplemental Table6 | Sequences of 48 crRNAs targeting H1N1 PR8 IAV and non-targeting crRNAs**

**Supplemental Table 7 | Sequences of 22 predicted crRNAs targeting all coronaviruses.** The top 6 (PAC-MAN-T6) covering >90% of coronaviruses including all published SARS-CoV-2 strains, SARS-CoV, and MERS-CoV are highlighted.

**Supplemental Table 8 | The crRNA-N18f can target all 202 sequenced SARS-CoV-2 genomes, including S and L strains.** Different strains of SARS-CoV-2 are shown in labels on the right: S, L, or unknown (U). * indicates the SARS-CoV-2 sequence that carries a single mutation in the crRNA-N18f targeting region.
