## Supplemental Table 5 for "Development of CRISPR as a prophylactic strategy to combat novel coronavirus and influenza"

Table S5

All Segment 1 Length 2341 vRNA

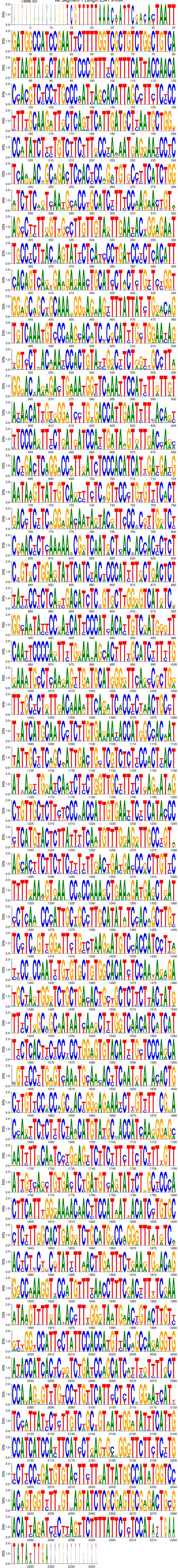

WebLogo 3.6.0

All Segment 2 Length 2341 vRNA

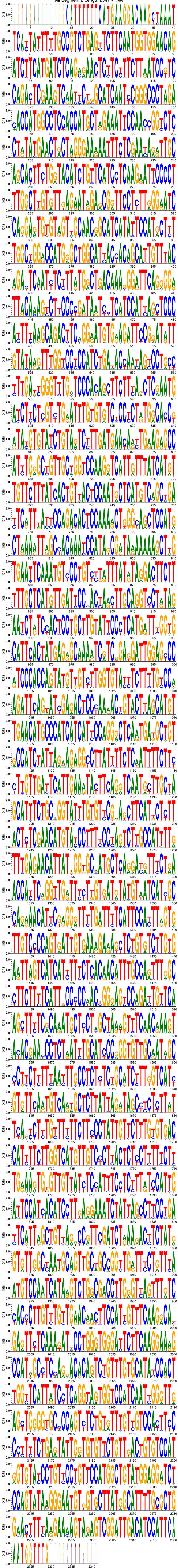

All Segment 3 Length 2234 vRNA

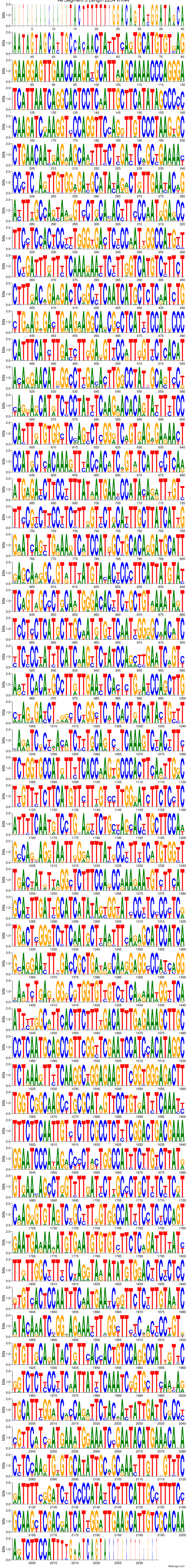

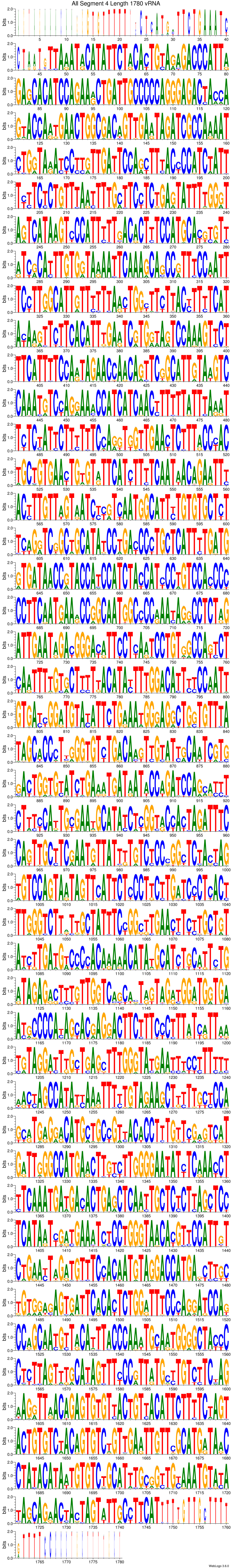
All Segment 4 Length 1780 vRNA
bits
WebLogo 3.6.0

All Segment 5 Length 1567 vRNA

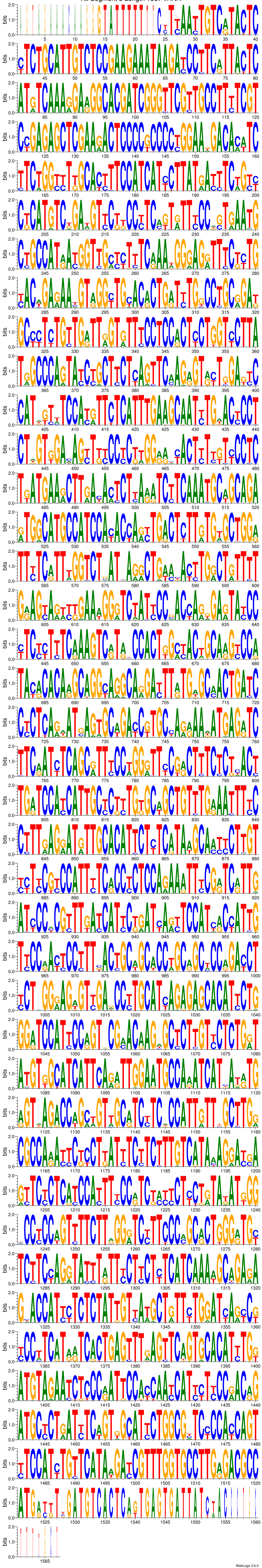

All Segment 6 Length 1477 vRNA

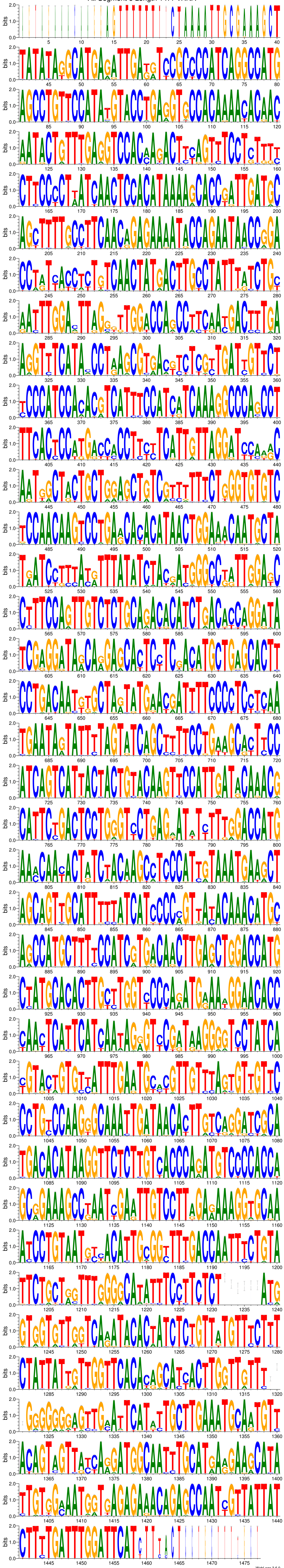

WebLogo 3.6.0

### All Segment 7 Length 1027 vRNA

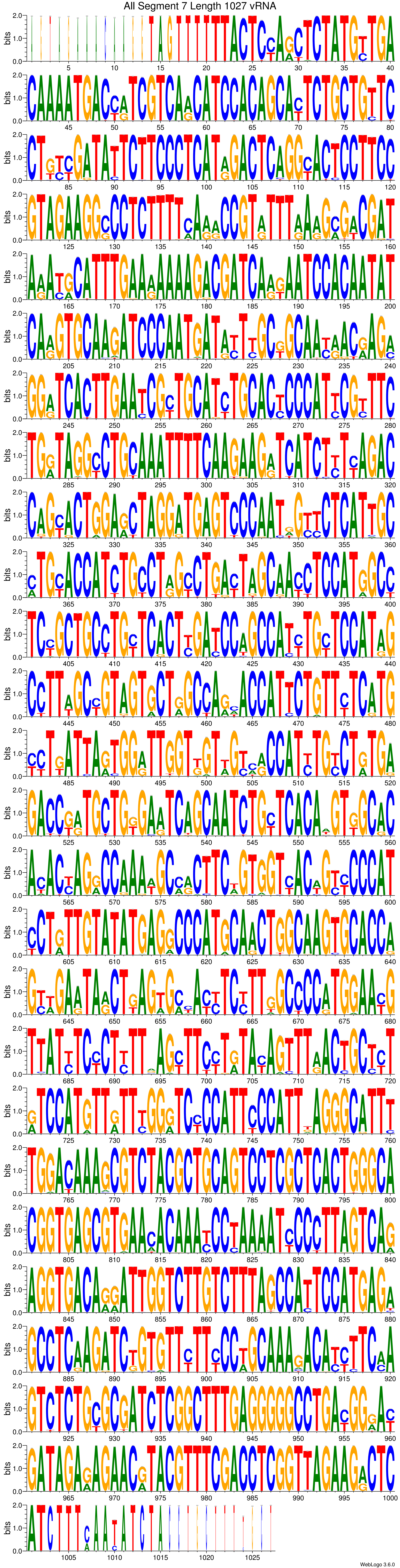

All Segment 8 Length 890 vRNA

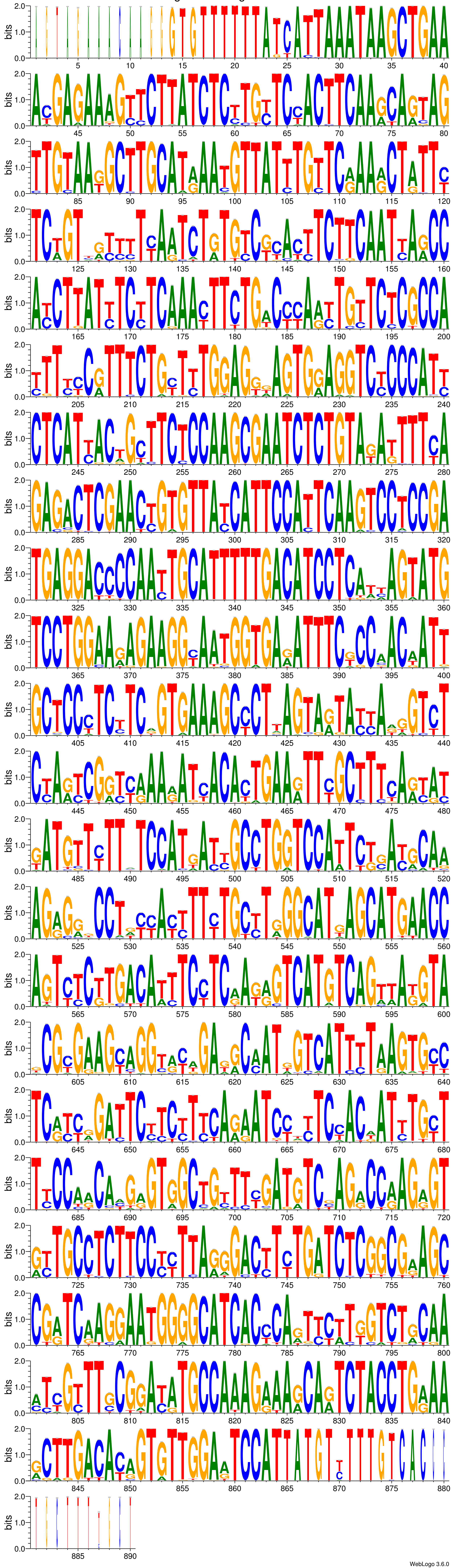
