## Supplemental Table 8 for "Development of CRISPR as a prophylactic strategy to combat novel coronavirus and influenza"

### Supplemental Table 8 l The tested crRNA can target 202 SARS-CoV-2 including S and L strains

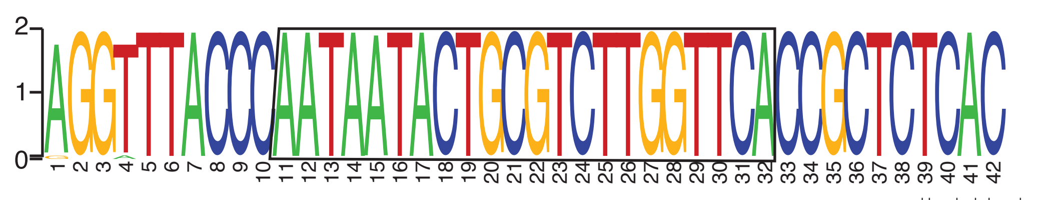


28,800 **crRNA-N18f** 3' UUAUUAUGACGCAGAACCAAGU 5' 28,841

| Sequence | Alignment | S/L/U strains |
|---|---|---|
| Consensus | 5' AGGTTTACCCAATAATACTGCGTCTTGGTTCACCGCTCTCAC 3' | |
| Identity | | |
| 1. EPI_ISL_402119 | AGGTTTACCCAATAATACTGCGTCTTGGTTCACCGCTCTCAC | L |
| 2. EPI_ISL_402121 | AGGTTTACCCAATAATACTGCGTCTTGGTTCACCGCTCTCAC | L |
| 3. EPI_ISL_402120 | AGGTTTACCCAATAATACTGCGTCTTGGTTCACCGCTCTCAC | L |
| 4. EPI_ISL_402123 | AGGTTTACCCAATAATACTGCGTCTTGGTTCACCGCTCTCAC | L |
| 5. EPI_ISL_402124 | AGGTTTACCCAATAATACTGCGTCTTGGTTCACCGCTCTCAC | L |
| 6. EPI_ISL_403962 | AGGTTTACCCAATAATACTGCGTCTTGGTTCACCGCTCTCAC | L |
| 7. EPI_ISL_403963 | AGGTTTACCCAATAATACTGCGTCTTGGTTCACCGCTCTCAC | L |
| 8. EPI_ISL_402127 | AGGTTTACCCAATAATACTGCGTCTTGGTTCACCGCTCTCAC | L |
| 9. EPI_ISL_402128 | AGGTTTACCCAATAATACTGCGTCTTGGTTCACCGCTCTCAC | L |
| 10. EPI_ISL_402129 | AGGTTTACCCAATAATACTGCGTCTTGGTTCACCGCTCTCAC | L |
| 11. EPI_ISL_402130 | AGGTTTACCCAATAATACTGCGTCTTGGTTCACCGCTCTCAC | S |
| 12. EPI_ISL_403932 | AGGTTTACCCAATAATACTGCGTCTTGGTTCACCGCTCTCAC | S |
| 13. EPI_ISL_403933 | AGGTTTACCCAATAATACTGCGTCTTGGTTCACCGCTCTCAC | L |
| 14. EPI_ISL_403934 | AGGTTTACCCAATAATACTGCGTCTTGGTTCACCGCTCTCAC | L |
| 15. EPI_ISL_403935 | AGGTTTACCCAATAATACTGCGTCTTGGTTCACCGCTCTCAC | S |
| 16. EPI_ISL_403936 | AGGTTTACCCAATAATACTGCGTCTTGGTTCACCGCTCTCAC | L |
| 17. EPI_ISL_403937 | AGGTTTACCCAATAATACTGCGTCTTGGTTCACCGCTCTCAC | L |
| 18. EPI_ISL_403930 | AGGTTTACCCAATAATACTGCGTCTTGGTTCACCGCTCTCAC | L |
| 19. EPI_ISL_402132 | AGGTTTACCCAATAATACTGCGTCTTGGTTCACCGCTCTCAC | L |
| 20. EPI_ISL_404227 | AGGTTTACCCAATAATACTGCGTCTTGGTTCACCGCTCTCAC | L |
| 21. EPI_ISL_404228 | AGGTTTACCCAATAATACTGCGTCTTGGTTCACCGCTCTCAC | L |
| 22. EPI_ISL_402131 | AGGTTTACCCAATAATACTGCGTCTTGGTTCACCGCTCTCAC | U |
| 23. EPI_ISL_404895 | AGGTTTACCCAATAATACTGCGTCTTGGTTCACCGCTCTCAC | S |
| 24. EPI_ISL_407079 | AGGTTTACCCAATAATACTGCGTCTTGGTTCACCGCTCTCAC | U |
| 25. EPI_ISL_405839 | AGGTTTACCCAATAATACTGCGTCTTGGTTCACCGCTCTCAC | S |
| 26. EPI_ISL_406030 | AGGTTTACCCAATAATACTGCGTCTTGGTTCACCGCTCTCAC | S |
| 27. EPI_ISL_404253 | AGGTTTACCCAATAATACTGCGTCTTGGTTCACCGCTCTCAC | L+S |
| 28. EPI_ISL_406034 | AGGTTTACCCAATAATACTGCGTCTTGGTTCACCGCTCTCAC | S |
| 29. EPI_ISL_406036 | AGGTTTACCCAATAATACTGCGTCTTGGTTCACCGCTCTCAC | L |
| 30. EPI_ISL_406223 | AGGTTTACCCAATAATACTGCGTCTTGGTTCACCGCTCTCAC | S |
| 31. EPI_ISL_406531 | AGGTTTACCCAATAATACTGCGTCTTGGTTCACCGCTCTCAC | L |
| 32. EPI_ISL_406533 | AGGTTTACCCAATAATACTGCGTCTTGGTTCACCGCTCTCAC | L |
| 33. EPI_ISL_406534 | AGGTTTACCCAATAATACTGCGTCTTGGTTCACCGCTCTCAC | L |
| 34. EPI_ISL_406535 | AGGTTTACCCAATAATACTGCGTCTTGGTTCACCGCTCTCAC | L |
| 35. EPI_ISL_406536 | AGGTTTACCCAATAATACTGCGTCTTGGTTCACCGCTCTCAC | L |
| 36. EPI_ISL_406538 | AGGTTTACCCAATAATACTGCGTCTTGGTTCACCGCTCTCAC | L |
| 37. EPI_ISL_406592 | AGGTTTACCCAATAATACTGCGTCTTGGTTCACCGCTCTCAC | U |
| 38. EPI_ISL_406593 | AGGTTTACCCAATAATACTGCGTCTTGGTTCACCGCTCTCAC | S |
| 39. EPI_ISL_406594 | AGGTTTACCCAATAATACTGCGTCTTGGTTCACCGCTCTCAC | L |
| 40. EPI_ISL_406597 | AGGTTTACCCAATAATACTGCGTCTTGGTTCACCGCTCTCAC | L |
| 41. EPI_ISL_406596 | AGGTTTACCCAATAATACTGCGTCTTGGTTCACCGCTCTCAC | L |
| 42. EPI_ISL_406862 | AGGTTTACCCAATAATACTGCGTCTTGGTTCACCGCTCTCAC | L |
| 43. EPI_ISL_406844 | AGGTTTACCCAATAATACTGCGTCTTGGTTCACCGCTCTCAC | L |
| 44. EPI_ISL_406716 | AGGTTTACCCAATAATACTGCGTCTTGGTTCACCGCTCTCAC | L |
| 45. EPI_ISL_406717 | AGGTTTACCCAATAATACTGCGTCTTGGTTCACCGCTCTCAC | L |
| 46. EPI_ISL_406798 | AGGTTTACCCAATAATACTGCGTCTTGGTTCACCGCTCTCAC | L |
| 47. EPI_ISL_406800 | AGGTTTACCCAATAATACTGCGTCTTGGTTCACCGCTCTCAC | L |
| 48. EPI_ISL_406801 | AGGTTTACCCAATAATACTGCGTCTTGGTTCACCGCTCTCAC | S |
| 49. EPI_ISL_406970 | AGGTTTACCCAATAATACTGCGTCTTGGTTCACCGCTCTCAC | L |
| 50. EPI_ISL_406973 | AGGTTTACCCAATAATACTGCGTCTTGGTTCACCGCTCTCAC | L |
| 51. EPI_ISL_407073 | AGGTTTACCCAATAATACTGCGTCTTGGTTCACCGCTCTCAC | S |
| 52. EPI_ISL_407071 | AGGTTTACCCAATAATACTGCGTCTTGGTTCACCGCTCTCAC | S |
| 53. EPI_ISL_407214 | AGGTTTACCCAATAATACTGCGTCTTGGTTCACCGCTCTCAC | S |
| 54. EPI_ISL_407313 | AGGTTTACCCAATAATACTGCGTCTTGGTTCACCGCTCTCAC | L |
| 55. EPI_ISL_407893 | AGGTTTACCCAATAATACTGCGTCTTGGTTCACCGCTCTCAC | S |
| 56. EPI_ISL_407976 | AGGTTTACCCAATAATACTGCGTCTTGGTTCACCGCTCTCAC | S |
| 57. EPI_ISL_407988 | AGGTTTACCCAATAATACTGCGTCTTGGTTCACCGCTCTCAC | L |
| 58. EPI_ISL_407987 | AGGTTTACCCAATAATACTGCGTCTTGGTTCACCGCTCTCAC | L |
| 59. EPI_ISL_408010 | AGGTTTACCCAATAATACTGCGTCTTGGTTCACCGCTCTCAC | L |
| 60. EPI_ISL_408009 | AGGTTTACCCAATAATACTGCGTCTTGGTTCACCGCTCTCAC | L |
| 61. EPI_ISL_408008 | AGGTTTACCCAATAATACTGCGTCTTGGTTCACCGCTCTCAC | L |
| 62. EPI_ISL_408430 | AGGTTTACCCAATAATACTGCGTCTTGGTTCACCGCTCTCAC | L |
| 63. EPI_ISL_408431 | AGGTTTACCCAATAATACTGCGTCTTGGTTCACCGCTCTCAC | L |
| 64. EPI_ISL_408480 | AGGTTTACCCAATAATACTGCGTCTTGGTTCACCGCTCTCAC | S |
| 65. EPI_ISL_408481 | AGGTTTACCCAATAATACTGCGTCTTGGTTCACCGCTCTCAC | L |
| 66. EPI_ISL_408482 | AGGTTTACCCAATAATACTGCGTCTTGGTTCACCGCTCTCAC | L |
| 67. EPI_ISL_408484 | AGGTTTACCCAATAATACTGCGTCTTGGTTCACCGCTCTCAC | S |
| 68. EPI_ISL_408486 | AGGTTTACCCAATAATACTGCGTCTTGGTTCACCGCTCTCAC | L |
| 69. EPI_ISL_408488 | AGGTTTACCCAATAATACTGCGTCTTGGTTCACCGCTCTCAC | L |
| 70. EPI_ISL_408489 | AGGTTTACCCAATAATACTGCGTCTTGGTTCACCGCTCTCAC | S |
| 71. EPI_ISL_408514 | AGGTTTACCCAATAATACTGCGTCTTGGTTCACCGCTCTCAC | L |
| 72. EPI_ISL_408515 | AGGTTTACCCAATAATACTGCGTCTTGGTTCACCGCTCTCAC | L |
| 73. EPI_ISL_408665 | AGGTTTACCCAATAATACTGCGTCTTGGTTCACCGCTCTCAC | S |
| 74. EPI_ISL_408666 | AGGTTTACCCAATAATACTGCGTCTTGGTTCACCGCTCTCAC | S |
| 75. EPI_ISL_408667 | AGGTTTACCCAATAATACTGCGTCTTGGTTCACCGCTCTCAC | S |
| 76. EPI_ISL_408669 | AGGTTTACCCAATAATACTGCGTCTTGGTTCACCGCTCTCAC | L |
| 77. EPI_ISL_407084 | AGGTTTACCCAATAATACTGCGTCTTGGTTCACCGCTCTCAC | L |
| 78. EPI_ISL_403931 | AGGTTTACCCAATAATACTGCGTCTTGGTTCACCGCTCTCAC | L |
| 79. EPI_ISL_403929 | AGGTTTACCCAATAATACTGCGTCTTGGTTCACCGCTCTCAC | L |
| 80. EPI_ISL_403928 | AGGTTTACCCAATAATACTGCGTCTTGGTTCACCGCTCTCAC | L |
| 81. EPI_ISL_408670 | AGGTTTACCCAATAATACTGCGTCTTGGTTCACCGCTCTCAC | L |
| 82. EPI_ISL_406595 | AGGTTTACCCAATAATACTGCGTCTTGGTTCACCGCTCTCAC | L |
| 83. EPI_ISL_406031 | AGGTTTACCCAATAATACTGCGTCTTGGTTCACCGCTCTCAC | L |
| 84. EPI_ISL_408976 | AGGTTTACCCAATAATACTGCGTCTTGGTTCACCGCTCTCAC | L |
| 85. EPI_ISL_408977 | AGGTTTACCCAATAATACTGCGTCTTGGTTCACCGCTCTCAC | L |
| 86. EPI_ISL_410045 | AGGTTTACCCAATAATACTGCGTCTTGGTTCACCGCTCTCAC | S |
| 87. EPI_ISL_410044 | AGGTTTACCCAATAATACTGCGTCTTGGTTCACCGCTCTCAC | L |
| 88. EPI_ISL_409067 | AGGTTTACCCAATAATACTGCGTCTTGGTTCACCGCTCTCAC | L |
| 89. EPI_ISL_408478 | AGGTTTACCCAATAATACTGCGTCTTGGTTCACCGCTCTCAC | S |
| 90. EPI_ISL_408479 | AGGTTTACCCAATAATACTGCGTCTTGGTTCACCGCTCTCAC | L |
| 91. EPI_ISL_410218 | AGGTTTACCCAATAATACTGCGTCTTGGTTCACCGCTCTCAC | L |
| 92. EPI_ISL_408487 | AGGTTTACCCAATAATACTGCGTCTTGGTTCACCGCTCTCAC | U |
| 93. EPI_ISL_408483 | AGGTTTACCCAATAATACTGCGTCTTGGTTCACCGCTCTCAC | U |
| 94. EPI_ISL_408485 | AGGTTTACCCAATAATACTGCGTCTTGGTTCACCGCTCTCAC | U |
| 95. EPI_ISL_408978 | AGGTTTACCCAATAATACTGCGTCTTGGTTCACCGCTCTCAC | U |
| 96. EPI_ISL_410486 | AGGTTTACCCAATAATACTGCGTCTTGGTTCACCGCTCTCAC | U |
| 97. EPI_ISL_402125 | AGGTTTACCCAATAATACTGCGTCTTGGTTCACCGCTCTCAC | L |
| 98. EPI_ISL_410301 | AGGTTTACCCAATAATACTGCGTCTTGGTTCACCGCTCTCAC | L |
| 99. EPI_ISL_410532 | AGGTTTACCCAATAATACTGCGTCTTGGTTCACCGCTCTCAC | U |
| 100. EPI_ISL_410531 | AGGTTTACCCAATAATACTGCGTCTTGGTTCACCGCTCTCAC | U |
| 101. EPI_ISL_410535 | AGGTTTACCCAATAATACTGCGTCTTGGTTCACCGCTCTCAC | U |
| 102. EPI_ISL_410537 | AGGTTTACCCAATAATACTGCGTCTTGGTTCACCGCTCTCAC | U |
| 103. EPI_ISL_410536 | AGGTTTACCCAATAATACTGCGTCTTGGTTCACCGCTCTCAC | U |
| 104. EPI_ISL_407215 | AGGTTTACCCAATAATACTGCGTCTTGGTTCACCGCTCTCAC | S |
| 105. EPI_ISL_410539 | GGGATTACCCAATAATACTGCGTCTTGGTTCACCGCTCTCAC | U |
| * 106. EPI_ISL_410541 | GGGATTACCCAATAGTACTGCGTCTTGGTTCACCGCTCTCAC | U |
| 107. EPI_ISL_410540 | GGGATTACCCAATAATACTGCGTCTTGGTTCACCGCTCTCAC | U |
| 108. EPI_ISL_410538 | GGGATTACCCAATAATACTGCGTCTTGGTTCACCGCTCTCAC | U |
| 109. EPI_ISL_410543 | GGGATTACCCAATAATACTGCGTCTTGGTTCACCGCTCTCAC | U |
| 110. EPI_ISL_410542 | GGGATTACCCAATAATACTGCGTCTTGGTTCACCGCTCTCAC | U |
| 111. EPI_ISL_410544 | AGGTTTACCCAATAATACTGCGTCTTGGTTCACCGCTCTCAC | U |
| 112. EPI_ISL_410717 | AGGTTTACCCAATAATACTGCGTCTTGGTTCACCGCTCTCAC | U |
| 113. EPI_ISL_410718 | AGGTTTACCCAATAATACTGCGTCTTGGTTCACCGCTCTCAC | U |
| 114. EPI_ISL_410713 | AGGTTTACCCAATAATACTGCGTCTTGGTTCACCGCTCTCAC | U |
| 115. EPI_ISL_410716 | AGGTTTACCCAATAATACTGCGTCTTGGTTCACCGCTCTCAC | U |
| 116. EPI_ISL_407894 | AGGTTTACCCAATAATACTGCGTCTTGGTTCACCGCTCTCAC | S |
| 117. EPI_ISL_407896 | AGGTTTACCCAATAATACTGCGTCTTGGTTCACCGCTCTCAC | S |
| 118. EPI_ISL_410714 | AGGTTTACCCAATAATACTGCGTCTTGGTTCACCGCTCTCAC | U |
| 119. EPI_ISL_410715 | AGGTTTACCCAATAATACTGCGTCTTGGTTCACCGCTCTCAC | U |
| 120. EPI_ISL_410719 | AGGTTTACCCAATAATACTGCGTCTTGGTTCACCGCTCTCAC | U |
| 121. EPI_ISL_410545 | AGGTTTACCCAATAATACTGCGTCTTGGTTCACCGCTCTCAC | U |
| 122. EPI_ISL_410546 | AGGTTTACCCAATAATACTGCGTCTTGGTTCACCGCTCTCAC | U |
| 123. EPI_ISL_410720 | AGGTTTACCCAATAATACTGCGTCTTGGTTCACCGCTCTCAC | U |
| 124. EPI_ISL_410984 | AGGTTTACCCAATAATACTGCGTCTTGGTTCACCGCTCTCAC | U |
| 125. EPI_ISL_410721 | AGGTTTACCCAATAATACTGCGTCTTGGTTCACCGCTCTCAC | U |
| 126. EPI_ISL_411060 | AGGTTTACCCAATAATACTGCGTCTTGGTTCACCGCTCTCAC | U |
| 127. EPI_ISL_411066 | AGGTTTACCCAATAATACTGCGTCTTGGTTCACCGCTCTCAC | U |
| 128. EPI_ISL_411218 | AGGTTTACCCAATAATACTGCGTCTTGGTTCACCGCTCTCAC | U |
| 129. EPI_ISL_411219 | AGGTTTACCCAATAATACTGCGTCTTGGTTCACCGCTCTCAC | U |
| 130. EPI_ISL_411220 | AGGTTTACCCAATAATACTGCGTCTTGGTTCACCGCTCTCAC | U |
| 131. EPI_ISL_411902 | AGGTTTACCCAATAATACTGCGTCTTGGTTCACCGCTCTCAC | U |
| 132. EPI_ISL_411915 | AGGTTTACCCAATAATACTGCGTCTTGGTTCACCGCTCTCAC | U |
| 133. EPI_ISL_411926 | AGGTTTACCCAATAATACTGCGTCTTGGTTCACCGCTCTCAC | U |
| 134. EPI_ISL_411927 | AGGTTTACCCAATAATACTGCGTCTTGGTTCACCGCTCTCAC | U |
| 135. EPI_ISL_406799 | AGGTTTACCCAATAATACTGCGTCTTGGTTCACCGCTCTCAC | U |
| 136. EPI_ISL_407193 | AGGTTTACCCAATAATACTGCGTCTTGGTTCACCGCTCTCAC | S |
| 137. EPI_ISL_411950 | AGGTTTACCCAATAATACTGCGTCTTGGTTCACCGCTCTCAC | U |
| 138. EPI_ISL_411953 | AGGTTTACCCAATAATACTGCGTCTTGGTTCACCGCTCTCAC | U |
| 139. EPI_ISL_411952 | AGGTTTACCCAATAATACTGCGTCTTGGTTCACCGCTCTCAC | U |
| 140. EPI_ISL_411929 | AGGTTTACCCAATAATACTGCGTCTTGGTTCACCGCTCTCAC | L |
| 141. EPI_ISL_411954 | AGGTTTACCCAATAATACTGCGTCTTGGTTCACCGCTCTCAC | U |
| 142. EPI_ISL_411955 | AGGTTTACCCAATAATACTGCGTCTTGGTTCACCGCTCTCAC | U |
| 143. EPI_ISL_411956 | AGGTTTACCCAATAATACTGCGTCTTGGTTCACCGCTCTCAC | U |
| 144. EPI_ISL_411951 | AGGTTTACCCAATAATACTGCGTCTTGGTTCACCGCTCTCAC | U |
| 145. EPI_ISL_411957 | AGGTTTACCCAATAATACTGCGTCTTGGTTCACCGCTCTCAC | U |
| 146. EPI_ISL_412026 | AGGTTTACCCAATAATACTGCGTCTTGGTTCACCGCTCTCAC | U |
| 147. EPI_ISL_412028 | AGGTTTACCCAATAATACTGCGTCTTGGTTCACCGCTCTCAC | U |
| 148. EPI_ISL_412029 | AGGTTTACCCAATAATACTGCGTCTTGGTTCACCGCTCTCAC | U |
| 149. EPI_ISL_412030 | AGGTTTACCCAATAATACTGCGTCTTGGTTCACCGCTCTCAC | U |
| 150. EPI_ISL_408668 | AGGTTTACCCAATAATACTGCGTCTTGGTTCACCGCTCTCAC | S |
| 151. EPI_ISL_412459 | AGGTTTACCCAATAATACTGCGTCTTGGTTCACCGCTCTCAC | U |
| 152. EPI_ISL_412116 | AGGTTTACCCAATAATACTGCGTCTTGGTTCACCGCTCTCAC | U |
| 153. EPI_ISL_412862 | AGGTTTACCCAATAATACTGCGTCTTGGTTCACCGCTCTCAC | U |
| 154. EPI_ISL_412869 | AGGTTTACCCAATAATACTGCGTCTTGGTTCACCGCTCTCAC | U |
| 155. EPI_ISL_412870 | AGGTTTACCCAATAATACTGCGTCTTGGTTCACCGCTCTCAC | U |
| 156. EPI_ISL_412871 | AGGTTTACCCAATAATACTGCGTCTTGGTTCACCGCTCTCAC | U |
| 157. EPI_ISL_412872 | AGGTTTACCCAATAATACTGCGTCTTGGTTCACCGCTCTCAC | U |
| 158. EPI_ISL_412873 | AGGTTTACCCAATAATACTGCGTCTTGGTTCACCGCTCTCAC | U |
| 159. EPI_ISL_412898 | AGGTTTACCCAATAATACTGCGTCTTGGTTCACCGCTCTCAC | U |
| 160. EPI_ISL_412899 | AGGTTTACCCAATAATACTGCGTCTTGGTTCACCGCTCTCAC | U |
| 161. EPI_ISL_412900 | AGGTTTACCCAATAATACTGCGTCTTGGTTCACCGCTCTCAC | U |
| 162. EPI_ISL_412912 | AGGTTTACCCAATAATACTGCGTCTTGGTTCACCGCTCTCAC | U |
| 163. EPI_ISL_412964 | AGGTTTACCCAATAATACTGCGTCTTGGTTCACCGCTCTCAC | U |
| 164. EPI_ISL_412965 | AGGTTTACCCAATAATACTGCGTCTTGGTTCACCGCTCTCAC | U |
| 165. EPI_ISL_412966 | AGGTTTACCCAATAATACTGCGTCTTGGTTCACCGCTCTCAC | U |
| 166. EPI_ISL_412967 | AGGTTTACCCAATAATACTGCGTCTTGGTTCACCGCTCTCAC | U |
| 167. EPI_ISL_412968 | AGGTTTACCCAATAATACTGCGTCTTGGTTCACCGCTCTCAC | U |
| 168. EPI_ISL_412969 | AGGTTTACCCAATAATACTGCGTCTTGGTTCACCGCTCTCAC | U |
| 169. EPI_ISL_412971 | AGGTTTACCCAATAATACTGCGTCTTGGTTCACCGCTCTCAC | U |
| 170. EPI_ISL_412970 | AGGTTTACCCAATAATACTGCGTCTTGGTTCACCGCTCTCAC | U |
| 171. EPI_ISL_412973 | AGGTTTACCCAATAATACTGCGTCTTGGTTCACCGCTCTCAC | U |
| 172. EPI_ISL_412972 | AGGTTTACCCAATAATACTGCGTCTTGGTTCACCGCTCTCAC | U |
| 173. EPI_ISL_412974 | AGGTTTACCCAATAATACTGCGTCTTGGTTCACCGCTCTCAC | U |
| 174. EPI_ISL_412975 | AGGTTTACCCAATAATACTGCGTCTTGGTTCACCGCTCTCAC | U |
| 175. EPI_ISL_412978 | AGGTTTACCCAATAATACTGCGTCTTGGTTCACCGCTCTCAC | U |
| 176. EPI_ISL_412979 | AGGTTTACCCAATAATACTGCGTCTTGGTTCACCGCTCTCAC | U |
| 177. EPI_ISL_412980 | AGGTTTACCCAATAATACTGCGTCTTGGTTCACCGCTCTCAC | U |
| 178. EPI_ISL_412981 | AGGTTTACCCAATAATACTGCGTCTTGGTTCACCGCTCTCAC | U |
| 179. EPI_ISL_412982 | AGGTTTACCCAATAATACTGCGTCTTGGTTCACCGCTCTCAC | U |
| 180. EPI_ISL_412983 | AGGTTTACCCAATAATACTGCGTCTTGGTTCACCGCTCTCAC | U |
| 181. EPI_ISL_413016 | AGGTTTACCCAATAATACTGCGTCTTGGTTCACCGCTCTCAC | U |
| 182. EPI_ISL_413014 | AGGTTTACCCAATAATACTGCGTCTTGGTTCACCGCTCTCAC | U |
| 183. EPI_ISL_413017 | AGGTTTACCCAATAATACTGCGTCTTGGTTCACCGCTCTCAC | U |
| 184. EPI_ISL_413018 | AGGTTTACCCAATAATACTGCGTCTTGGTTCACCGCTCTCAC | U |
| 185. EPI_ISL_413019 | AGGTTTACCCAATAATACTGCGTCTTGGTTCACCGCTCTCAC | U |
| 186. EPI_ISL_413020 | AGGTTTACCCAATAATACTGCGTCTTGGTTCACCGCTCTCAC | U |
| 187. EPI_ISL_413021 | AGGTTTACCCAATAATACTGCGTCTTGGTTCACCGCTCTCAC | U |
| 188. EPI_ISL_413022 | AGGTTTACCCAATAATACTGCGTCTTGGTTCACCGCTCTCAC | U |
| 189. EPI_ISL_413023 | AGGTTTACCCAATAATACTGCGTCTTGGTTCACCGCTCTCAC | U |
| 190. EPI_ISL_413024 | AGGTTTACCCAATAATACTGCGTCTTGGTTCACCGCTCTCAC | U |
| 191. EPI_ISL_413213 | AGGTTTACCCAATAATACTGCGTCTTGGTTCACCGCTCTCAC | U |
| 192. EPI_ISL_413214 | AGGTTTACCCAATAATACTGCGTCTTGGTTCACCGCTCTCAC | U |
| 193. EPI_ISL_413221 | AGGTTTACCCAATAATACTGCGTCTTGGTTCACCGCTCTCAC | U |
| 194. EPI_ISL_413025 | AGGTTTACCCAATAATACTGCGTCTTGGTTCACCGCTCTCAC | U |
| 195. EPI_ISL_413455 | AGGTTTACCCAATAATACTGCGTCTTGGTTCACCGCTCTCAC | U |
| 196. EPI_ISL_413456 | AGGTTTACCCAATAATACTGCGTCTTGGTTCACCGCTCTCAC | U |
| 197. EPI_ISL_413015 | AGGTTTACCCAATAATACTGCGTCTTGGTTCACCGCTCTCAC | U |
| 198. EPI_ISL_413457 | AGGTTTACCCAATAATACTGCGTCTTGGTTCACCGCTCTCAC | U |
| 199. EPI_ISL_413458 | AGGTTTACCCAATAATACTGCGTCTTGGTTCACCGCTCTCAC | U |
| 200. EPI_ISL_413459 | AGGTTTACCCAATAATACTGCGTCTTGGTTCACCGCTCTCAC | U |
| 201. EPI_ISL_413485 | AGGTTTACCCAATAATACTGCGTCTTGGTTCACCGCTCTCAC | U |
| 202. EPI_ISL_413486 | AGGTTTACCCAATAATACTGCGTCTTGGTTCACCGCTCTCAC | U |
